# Gradual entry into carbon starvation decreases the death rate of *Escherichia coli*

**DOI:** 10.1101/2024.07.18.604087

**Authors:** Rossana Droghetti, Zara Gough, Hamid Seyed-Allaei, Severin Josef Schink, Ulrich Gerland

## Abstract

Bacterial fitness is determined both by how fast cells grow in nutrient-rich environments and by how well they survive when nutrients are depleted. However, these behaviors are not independent, since the molecular composition of non-growing cells is affected by their prior growth history. For instance, recent work observed that the death rates of *Escherichia coli* cultures that rapidly entered carbon starvation depend on their prior growth rates, with faster growth leading to exponentially faster death. On the other hand, it is well known that cells adapt their molecular composition as they slow down growth and enter stationary phase, which is generally believed to improve their chance of survival. Hence, the question arises to what extent this adaptation process reduces the subsequent death rate. And how does the duration of the time window during which cells are allowed to adapt determine the reduction in death rate, and thus the fitness benefit of adaptation? Here, we study these quantitative questions by probing the adaptation of *E. coli* during gradual transitions from exponential growth to carbon starvation. We monitor such transitions in cultures with different initial growth conditions and measure the resulting rates of cell death after the transition. Our experiments demonstrate that cells with the opportunity to adapt their proteome composition before entering a state of starvation exhibit lower death rates compared to those that cannot, across various substrate conditions. The quantitative data is consistent with a theoretical model built on the assumption that before starvation, cells up-regulate a specific sector of the proteome, the effect of which is to decrease the death rate in energy-limiting conditions. This work highlights the influence of the non-genetic memory of a cell, specifically in the form of inherited proteome composition, on bacterial fitness. Our results emphasize that a comprehensive understanding of bacterial fitness requires quantitative characterization of bacterial physiology in all phases of their life cycle, including growth, stationary phase, and death, as well as the transitions between them.

**AUTHOR SUMMARY:** Bacteria inhabit dynamic environments and are frequently challenged by scarcity of nutrients. A recent study uncovered a curious link – faster bacterial growth leads to more rapid death when resources run out. We find that bacteria that gradually enter starvation exhibit significantly enhanced survival compared to those that do not have the chance to adapt. We interpret the observed quantitative behavior with the help of a theoretical model, which shows that our data is not compatible with a passive adaptation process, which would rely only on the general remodeling of the cellular proteome that is associated with growth transitions. Instead, our data are consistent with an active adaptation via up-regulation of genes that enhance survival during starvation. These results provide a novel perspective on bacterial survival strategies and underscore the importance of quantitatively investigating all phases of bacterial life cycles.

## INTRODUCTION

Both commensal and pathogenic bacteria have biphasic lifestyles, with a host-dependent, nutrient-rich phase, and a host-independent, nutrient-poor phase [1]. The fitness of these bacteria, i.e., their long-term reproductive success, then depends both on their ability to grow in the nutrient-rich phases and their ability to survive in the nutrient-poor phases. However, these abilities are not independent. For *E. coli*, it was recently shown that the rate of cell death in carbon-starved cell culture is substantially higher when the cells came from a rapidly growing culture, compared to when they came from a slow-growing condition [2]. A possible explanation of this history-dependent behavior is that the cell death rate is determined by the cell’s proteome composition, i.e. the set of all the proteins in a cell at a certain time. This composition is linked to the cell’s growth rate via well-established empirical growth laws [3]. According to this hypothesis, bacteria will enter starvation with distinct compositions of their proteome depending on the prior growth condition and the length of the adaptation phase, and the final proteome composition will then determine the death rate. This hypothesis potentially explains the exponential relation between growth and death rate found in ref. [2] (because the proteome changes with growth rate [3, 4]), and is consistent with mechanisms of non-genetic memory in *E. coli* [5, 6].

In rapidly fluctuating environments, non-genetic memory can be advantageous for bacteria, in particular when the environment alternates between two different carbon sources [5]. In contrast, when fast growth alternates with long periods of carbon starvation, the memory of previous rapid growth is detrimental because it leads to more rapid cell death during famine [2]. To alleviate this detrimental effect, cells adapt to carbon starvation, during (and after) the transition to the nutrient-poor environment. Much is known about the molecular processes governing the adaptation to carbon starvation [1, 7], but the resulting physiological behavior has not yet been characterized quantitatively, and it is still not well understood to what degree non-genetic memory impacts the adaptation of cells and, therefore, their long-term fitness. Here, we test the proteome-memory hypothesis by studying this adaptation behavior, in particular by testing the speed and degree of adaptation during the transition to carbon starvation.

To test the link between death rate and proteome composition hypothesized in ref. [2], one needs to understand how the composition changes during the shift towards starvation: the authors extrapolate a quantitative model for the kinetics of growth transitions [8] to situations in which the external carbon source is fully depleted by the growing cells. The model, called Flux-Controlled Regulation or FCR model, describes the dynamic remodeling of the proteome that occurs when the external condition changes and, as a response, cells change the regulation of protein synthesis of the different proteome sectors. Specifically, this coarse-grained kinetic model incorporates the regulation of the carbon catabolism and ribosomal proteome sectors, in a way such that the well-established steady-state growth laws are reproduced [3, 9]. The FCR model is quantitatively consistent with a wide range of growth transitions between different carbon sources, without invoking any protein degradation. Hence, the old proteome associated with conditions before the growth transition is primarily diluted by growth, such that it approaches the steady-state proteome composition associated with the new environment on a timescale that is inversely proportional to the growth rate. This “proteome memory” could limit the speed and degree of the adaptation to carbon starvation, since slow growth in carbon-poor conditions requires a long time to “delete the memory” of previous rich carbon conditions.

Here, we refer to the extrapolation of the FCR kinetic model for growth transitions to the entry into carbon starvation as the “proteome memory model”. It represents a drastic simplification of the actual adaptation process in *E. coli*, albeit a useful one: comparing it to the actual adaptation dynamics reveals the effects of additional regulatory mechanisms beyond the global, growth-dependent proteome remodeling predicted by the FCR model. Such a comparison is our central aim here. Towards this end, we perform experiments in which bacterial cultures enter carbon starvation in different ways, such that the transition ranges from abrupt to very gradual (Fig. 1). As expected, we observe that gradual entry into carbon starvation lowers the subsequent rate of cell death in the culture. However, this adaptation is stronger than predicted by the proteome memory model. To get some insight into the physiological nature of this adaptation, we make two hypotheses: that the cell during the transition to starvation is either (I) actively up-regulating a set of proteins that improves starvation survival, or (II) actively down-regulating another set of proteins that is detrimental for survival, for example, because they consume energy. We compared these two different models of additional targeted regulation with the experimental data we collected, and we found only the survival sector up-regulation one to be consistent with the data.

**FIG. 1:**
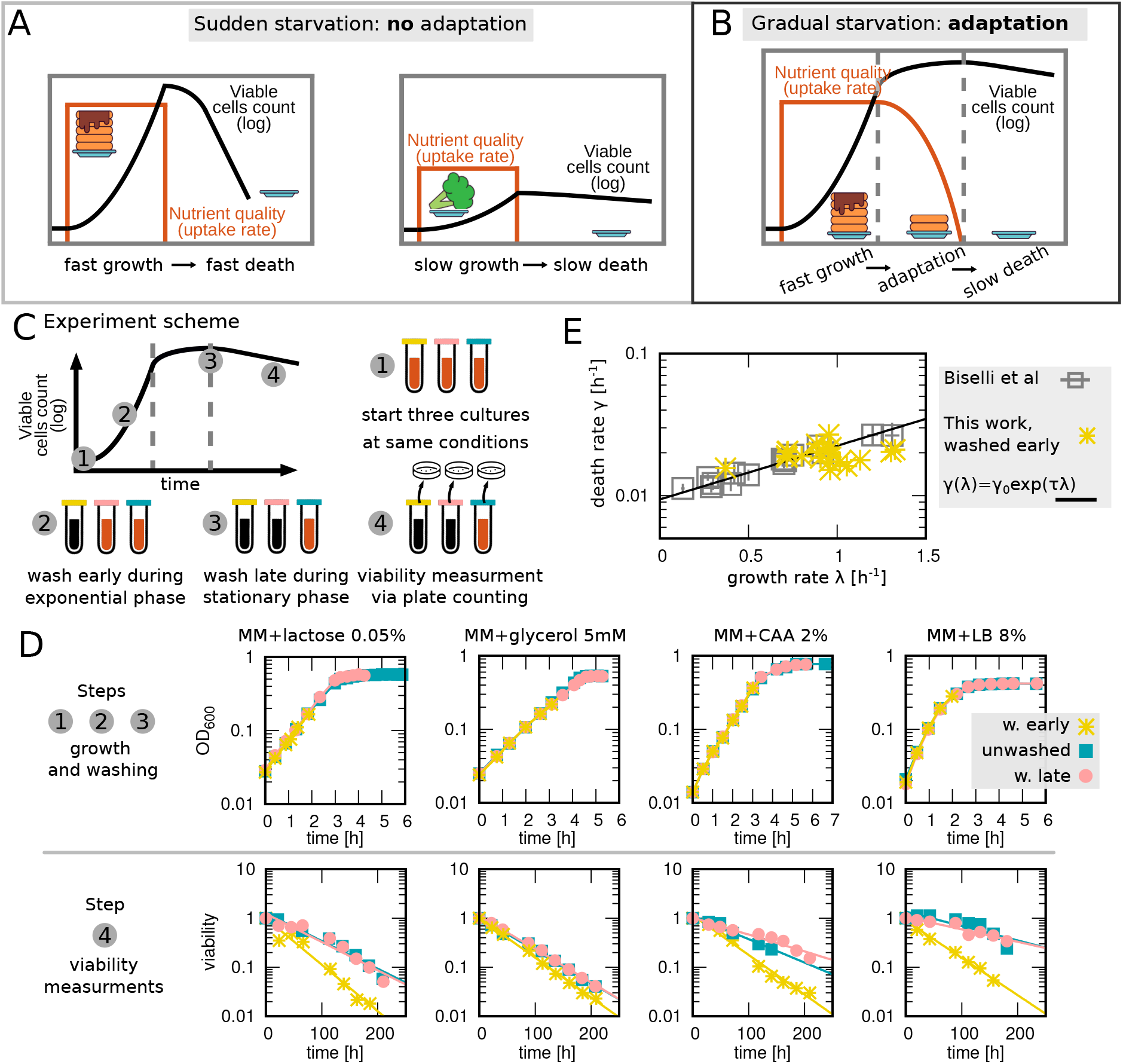
Bacterial adaptation to starvation during entry of stationary phase. (A) Graphical synopsis of the starting point of this study: when cells are suddenly carbon-starved after exponential growth, they die more rapidly the faster they previously grew [2]. (B) Graphical synopsis of adaptation during the entry into starvation: Cells experiencing a gradual entry into starvation can sense the carbon limitation and adapt to survive longer. (C) Experimental design: (I) Prepare three identical cultures and grow them at a constant temperature; (II) During exponential growth, one culture is washed (extracted by centrifugation and re-suspended in carbon-free medium), such that it abruptly enters stationary phase; (III) once the other cultures have reached stationary phase, one of them is washed to prevent potential further adaptation on leftover nutrients or metabolic byproducts; (IV) measure the cultures survival dynamics via plate counting. Bacterial adaptation to starvation during entry of stationary phase. (A) Graphical synopsis of the starting point of this study: when cells are suddenly carbon-starved after exponential growth, they die more rapidly the faster they previously grew [2]. (B) Graphical synopsis of adaptation during the entry into starvation: Cells experiencing a gradual entry into starvation can sense the carbon limitation and adapt to survive longer. (C) Experimental design: (I) Prepare three identical cultures and grow them at a constant temperature; (II) During exponential growth, one culture is washed (extracted by centrifugation and re-suspended in carbon-free medium), such that it abruptly enters stationary phase; (III) once the other cultures have reached stationary phase, one of them is washed to prevent potential further adaptation on leftover nutrients or metabolic byproducts; (IV) measure the cultures survival dynamics via plate counting. (D) Experiments show the impact of adaptation on viability. All the panels show data from this study, gathered for the three different cultures growing on the same substrate (e.g. MM+Lactose 0.5 % in the first column), “w. early” (yellow asterisks) is washed during exponential growth, while “w. late” (pink circles) is washed during the stationary phase. Blue squares represent unwashed cultures. The top row shows the first three steps of the experiment, where growth dynamics is measured in different mediums, and each point is an OD_600_ measurement. The bottom row shows viability during starvation. Each point is a viability measurement, the y coordinate shows the residual viability with respect to the start of starvation and the x coordinate is the time from starvation start. The plots show that the death phase is exponential for all of the cultures in all the tested mediums and that the “w. early” culture dies at a higher rate. (E) Our results for non-adapted (“washed early”) cultures are compatible with ref. [2]. Points are experimental results from Ref. [2] (grey squares) and this study (yellow asterisks). For each point, the x coordinate is the exponential growth rate, while the y one is the death rate. The black line shows the exponential growth-death relationship of Ref. [2] (see main text for parameter values). The panel shows that our results from non-adapted (washed early) cultures are compatible with the previously published data. For detailed experimental procedures, see “Materials and Methods”.

## RESULTS

### Gradual entry into carbon starvation reduces the death rate of *E. coli* cultures

We first quantify the extent to which cells adapt during gradual carbon starvation, by measuring how much the death rate of bacteria is reduced when they enter carbon starvation gradually rather than suddenly (Fig. 1A-B). Specifically, we grew several identical cultures of a wild-type *E. coli* strain (NCM3722) on a carbon-limited medium, and measured their growth rates *λ* in mid-exponential phase. We prevented the adaptation of some cultures by washing them with a carbon-free medium during exponential growth (“washed early”), while letting the other cultures adapt to stationary phase by nutrient depletion (Fig. 1C, see ‘Materials and Methods’ for details). We then either washed the adapted cultures shortly after growth had halted (“washed late”) to prevent further potential adaptation on leftover nutrients or metabolic byproducts, or skipped the washing step entirely (“unwashed”). Finally, we measured the decrease of the viability of all cultures over several days, by plating samples from the cultures, which were kept at a constant temperature of 37^°^C in a water bath shaker. Here, viability is defined as the number of colony-forming units (CFU) per milliliter (ml) of culture, and we plot the relative viability with respect to the starting time of starvation on a logarithmic y-axis. Fig. 1D (bottom) shows four examples of cultures grown on different media.

For all cultures, i.e., both the non-adapted (“washed early”) and the adapted (“washed late”, “unwashed”) cultures, the viability curves were consistent with an exponential decrease, such that an effective death rate *γ* could be extracted by a linear fit to the log-scale data. Fig. 1E shows that our death rates *γ* for the non-adapted (“washed early”) cultures are consistent with the empirical growth-death rate relation of Ref. [2, 10],

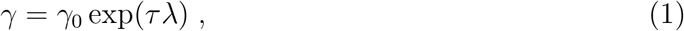

where *λ* is the prior growth rate of the culture, *γ*_0_ = 0.23*/*day represents the extrapolated death rate for a zero prior growth rate, while *τ* = 0.87 h is the timescale associated with the exponential increase of the death rate with the prior growth rate (corresponding to the slope in the semi-logarithmic plot of Fig. 1E).

Compared to the non-adapted cultures, the respective adapted cultures displayed a death rate that was up to three-fold lower. For instance, for the case of lactose as the carbon source, we obtained a death rate of 0.26 *±* 0.04/day for the “unwashed” cultures, whereas the non-adapted (“washed early”) death rate was 0.50 *±* 0.03/day (mean and standard deviation calculated from 7 replicates each). In contrast, there was little difference in death rate between “washed late” and “unwashed” cultures. In the lactose example, we obtained a death rate of 0.29 *±* 0.05/day for the “washed late” cultures, not significantly different from the 0.26 *±* 0.04/day rate for the “unwashed” cultures (p-value 0.151 for Welch’s t-test). Together, these observations suggest that the adaptation primarily occurs during the entry to stationary phase, rather than during stationary phase itself.

We observed different degrees of adaptation to starvation, depending on the medium used for the prior growth phase (Fig. 1D, Supplementary Figs. S2–S6, and Supplementary Table S1). The highest degrees of adaptation were produced by our complex growth media, casamino acids (CA) and lysogeny broth (LB). Growth of *E. coli* on these media displays a prolonged retardation phase, from exponential growth at a maximal rate to zero growth in stationary phase, as the different carbon sources contained within the media run out sequentially. During this retardation phase, cells can adapt to carbon starvation. Considering minimal media with a single carbon source, we observed a much stronger adaptation effect for lactose than for glycerol. One factor contributing to this difference may be that *E. coli* has a low affinity for lactose, with a Monod constant [11]) of *K*_*M*_ = 400*µM* [12], while the affinity for glycerol is high, with *K*_*M*_ = 5.6*µM* [13]. The Monod constant sets the extracellular concentration of the carbon source, at which the growth rate is reduced to half of the maximal growth rate on this carbon source. Hence, for a larger *K*_*M*_, the cell perceives the nutrient limitation at a higher concentration, and can use a larger pool of remaining external nutrient to remodel its molecular content for adaptation to starvation. Another factor contributing to this difference may be overflow metabolism [14]: cells growing on lactose excrete acetate, whereas those growing on glycerol do not. The excreted acetate may fuel adaptation to starvation. However, our observation that “washed late” and “unwashed” cultures behave similarly suggests that this factor is less significant. Finally, minimal medium with glucose as a carbon source produces a significant adaptation effect (Fig. S4) despite the high affinity, *K*_*M*_ = 5*µM*, for glucose [15]. This difference to the glycerol case may arise from a combination of factors, including acetate excretion and the different growth dynamics during the transition to stationary phase.

Taken together, the data suggests that the degree of adaptation depends in a complex way on the growth dynamics during the retardation phase between balanced exponential growth and the entry into stationary phase. In order to analyze this relationship, and to interpret our data quantitatively, we turn to a mathematical model for the adaptation process.

### Rationale of the model

Several regulatory genes have been identified as essential during starvation [17–21], above all the general stress response regulator *rpoS* [7, 22], but it is still unclear to what extent they are limiting for survival [23]. Moreover, previous experiments [2] found that *rpoS* mutants still display the growth-death relation (1), only with a different slope. Hence, as *rpoS* -mutant cells can still exhibit different degrees of starvation survival, implying that the existing genetic knowledge is incomplete.

The dependence of the death rate on the prior growth history observed in Fig. 1 is a form of non-genetic memory of cells. A likely candidate for the source of this memory is the composition of the cell’s proteome, which is history-dependent and known to change only slowly during growth transitions [8]. We therefore base our model on the assumption that the proteome composition at the entry of carbon starvation determines the subsequent death rate. Depending on the prior growth conditions and the length of the adaptation phase, cells will enter starvation with distinct compositions of their proteome, which then determine the death rate. This could simultaneously explain the growth-death relationship (1) (because the proteome changes with growth rate [3, 4]) and how adaptation decreases death rate (because the proteome changes during nutrient shifts [8] and exhaustion [16]). Cultures that were washed early during the exponential phase would then die faster, because they were unable to adapt their proteome.

Given that it is not clear which specific proteins are setting the death rate, we define the death rate as a general function of the proteome composition, a choice made also in previous work [24]. Because the majority of proteins follow distinct, collective patterns during exponential growth [3, 4] and adaptation [8], we assume that the death rate can be expressed as a function of all proteome sectors *{ϕ*_i_*}*,

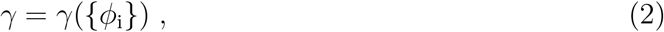

where each sector *ϕ*_i_ includes a set of co-regulated proteins that are either beneficial for survival (a “survival sector” *ϕ*_S_), or detrimental for survival (a “harmful sector” *ϕ*_H_). The size of these sectors would then determine the death rate. Note that *ϕ*_H_ will not be harmful in general, but only specifically during starvation. Examples of beneficial proteins could be proteins that scavenge reactive oxygen species [25–27], that protect DNA [21] or improve cell envelope integrity [23]. Harmful proteins could include those that waste energy or nutrients by futile cycling, e.g. unnecessary pumping or metabolic reactions. A priori, it is unclear which of the two sectors is the dominant contributor to changes in the death rate. In principle, cells can implement two adaptation strategies to lower the death rate: upregulate *ϕ*_S_ or down-regulate *ϕ*_H_, as illustrated in Fig. 2B.

**FIG. 2:**
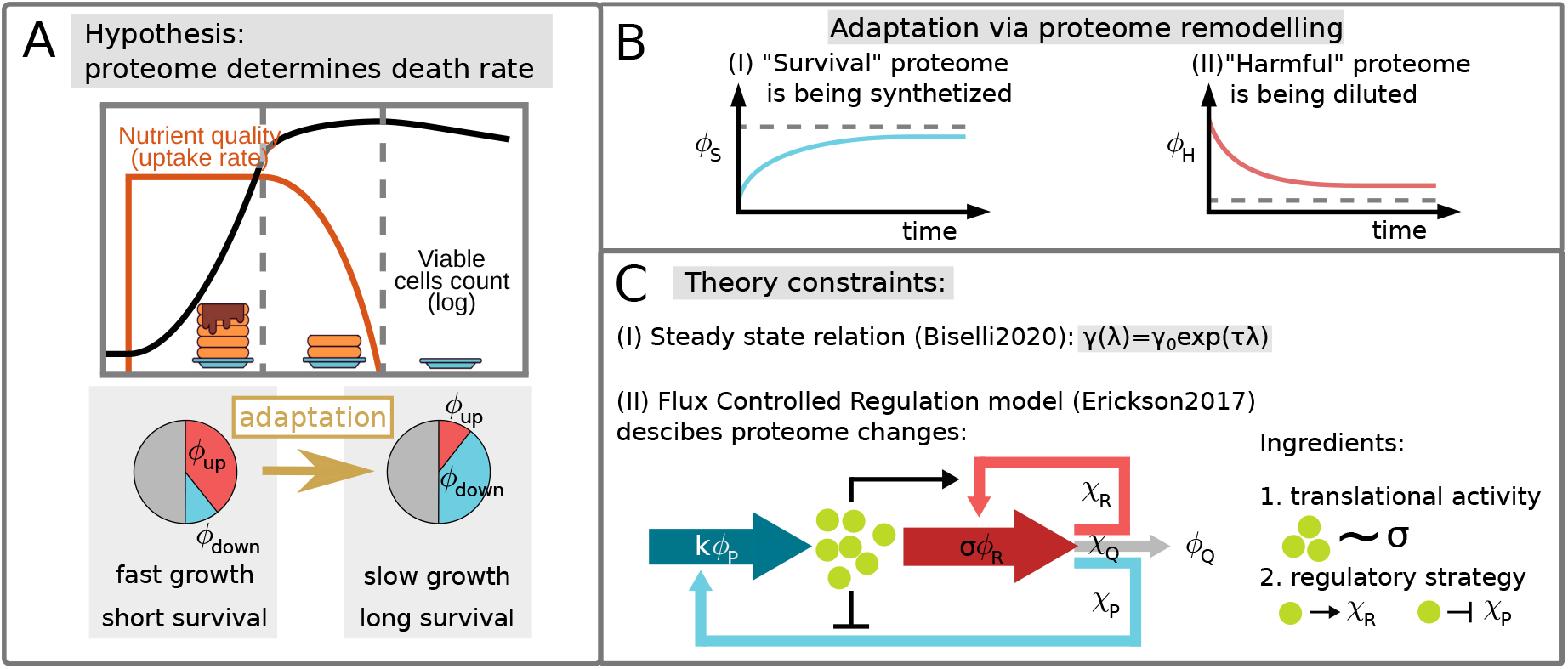
Rationale of the model. (A) Graphical synopsis of the hypothesis underlying our model, which is that the proteome composition at the entry of carbon starvation determines the subsequent death rate. During growth transitions, the proteome composition of a cell changes [8, 16], such that cells that have a smooth entry into starvation have a different composition compared to the composition during balanced exponential growth. In our model, the difference in death rates between adapted and non-adapted cultures is linked to this changed composition. (B) Cells can adapt to carbon starvation by (i) upregulating a “survival” proteome sector *ϕ*_S_, or (ii) down-regulating a “harmful” proteome sector *ϕ*_H_ that is not harmful in general, but only during starvation. (C) Our model takes into account two constraints: (i) it needs to reproduce the relation (1) between the steady-state growth and death rates [2]; (ii) during the adaptation phase cells cannot allocate their resources arbitrarily, but are constrained by the nutrient influx and the reaction of the regulatory circuits. The cellular resource allocation strategy in out-of-steady-state scenarios is described by the Flux Controlled Regulation (FCR) model of Ref. [8]. The main assumptions of this model is that the metabolic state of the cell (pool of central precursors) is reflected by the ribosome translational activity *σ*, defined as the average elongation rate. On a molecular level, the state of the precursor pool regulates the synthesis of the different proteome sectors via the regulatory functions 𝒳_up_ and 𝒳_down_.

### Proteome memory model for proteome-composition dependent death rate

In this section, we mathematically express the death rate in terms of proteome composition, *γ*(*{ϕ*_i_*}*), resulting in a model that we refer to as the “proteome memory model”. Our starting point is the relation (1) between *λ* and *γ*. From that equation, we derive a relation between *γ* and either the survival *ϕ*_S_ or the harmful sector *ϕ*_H_. During steady-state growth, growth rate and the proteome composition are strictly related [3, 4], with the exact relation depending on the type of growth limitation applied: for carbon limitation, which is the condition we are exploring in this work, the growth rate is related to nutrient quality. For other limitations, additional factors can affect the growth rate [4].

For each growth limitation, one can define three sectors: one that increases as the growth rate increases, one that decreases as the growth rate increases, and the one that stays constant [4]. These three classes enclose all the proteins, even if the slope at which their abundance changes is protein-specific. For growth in carbon-limited substrates, the up-regulated sector *ϕ*_up_ includes ribosomal proteins and many metabolic genes [3, 4]. The down-regulated sector *ϕ*_down_, on the other hand, includes carbon transporters, locomotion, and stress response genes [4, 28, 29]. What about the survival and harmful proteome? Given that high growth rates are associated with high death rates (for growth in carbon limited substrates [2, 23]) we can conclude that the survival sector would be one of the “down” sectors, and the harmful one of the “up”.

We make the hypothesis that these “up” and “down” sectors are a linear function of the growth rate (like the vast majority of all defined sectors [4]). This hypothesis allows us to write a growth law for the up and down sectors, which can be written as a function of a particular steady-state configuration (*λ*^*^ 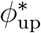) and (*λ*^*^,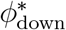) that can be interpreted as the steady state of the non-adapted (washed early) culture. Please refer to the Methods section for all the steps. By inserting the growth laws into Eq. (1), we obtain two functions that link the size of these sectors to the death rate:

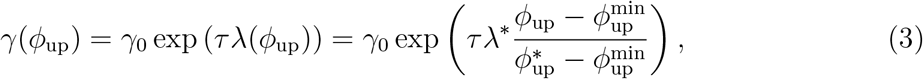

and

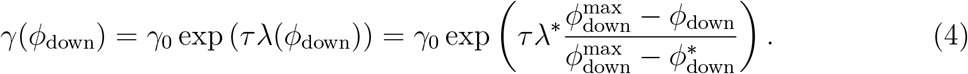

These two equations mathematically represent our proteome memory model, and they are central in the comparison between model and experimental results. We call the two terms inside the parenthesis 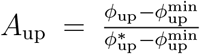 and 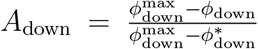 “adaptation extend”, as they represent how much the proteome has adapted from the exponential growth steady state compositions (*λ*^*^,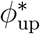) and (*λ*^*^, 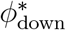), and quantifies the difference, in term of proteome composition, between the non adapted and adapted cultures. It is easy to see that if the culture is not able to adapt, then sectors will have a size that is equal to the steady state value: *ϕ*_up_ = 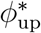 and *ϕ*_down_ = 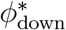. By putting these values into *A*_up_ and *A*_down_ we obtain *A*_up_ = *A*_down_ = 1, which leads to the same death rate of the non-adapted culture: *γ*_0_ exp (*τλ*^*^). As the cells adapt their proteome composition, the value of *A* parts from 1, resulting in a lower predicted death rate.

If we assume a model where the protein production is tightly related to the substrate availability, which provides nutrients that are converted into biomass, we can expect that for cultures washed early the sudden entry to the stationary phase means that they cannot adapt their proteome for starvation survival, and the proteome composition is completely determined by their pre-starvation growth rate. For adapted cultures, on the other hand, the proteome composition will change depending on the details of the entry to the stationary phase. To understand how much cells can adapt their proteome, we next turn to a kinetic model of protein regulation during the entry to the stationary phase.

#### Flux-controlled regulation model

The flux-controlled regulation (FCR) model [8] can describe the growth dynamics when the external environment is not constant, for example during nutrient up- and downshifts, based on the assumption that the proteome is regulated in a global, sector-defined fashion. In the model, a change in nutrient conditions, for example, a shift or nutrient depletion, will cause a change in the availability of the “central precursors”, i.e. the building blocks needed for the biosynthesis of new proteins. In particular, when the substrate is running out the amount of nutrients that the cell can gather from the environment becomes lower, and therefore the synthesis of new amino acids and other precursors needed for protein synthesis is slowed down. As a consequence of this, the translational activity *σ*, which is the average speed at which biosynthesis is carried out by ribosomes, starts to drop [8, 30]. The cell senses the change in *σ* via the ppGpp circuit [31] and adjusts the allocation of resources towards each sector accordingly, which is implemented in the model by making the regulatory functions χ_i_ a function of the translational activity, χ_i_ (*σ*). This allows the FCR model to describe how dynamic changes in the allocation of resources affect proteome composition and growth.

In the FCR model, the size of the sectors is determined by the sector-specific regulatory functions χ_i_, which are defined as the portion of the total biosynthesis flux redirected towards sector *i*,

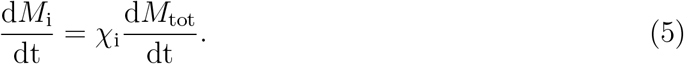

where *M*_i_ represents the mass of the *i*-th sector and *M*_tot_ the total biomass. From this mass equation, we can derive the differential equation for the sector size, defined as the ratio of the mass of the sector to the total protein mass 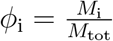,

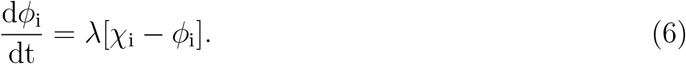

When *σ* drops during the last phase of growth, cells immediately adjust the regulatory functions *χ*_i_ accordingly. But because χ_i_ only affects the derivative of *ϕ*_i_, the size of the sectors adapts only gradually, and this adjustment is particularly slow when the growth rate is decreasing like in the shifts towards starvation that we are studying because the change in sector size is proportional to growth rate *λ*, Eq. 6. This model describes protein regulation strategy assuming that all the sectors are regulated in a global way because the kinetic of all the regulatory functions *χ*_i_ is given by the translational activity *σ*, therefore all of the sectors adapt following the same time-dependent kinetics.

Next, we need to understand how the survival and harmful sectors are regulated. We have hypothesize that *ϕ*_H_ ∈ *ϕ*_up_ and *ϕ*_S_ ∈ *ϕ*_down_, and now we need to define their regulatory functions. The FCR model assumes that the relation χ(*σ*) is in a quasi-steady state during growth transitions, meaning that the relation found in steady state growth conditions holds even during the adaptation phase, which translates into

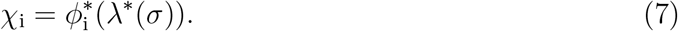

where the asterisks denote relations between *ϕ*_i_ and *λ* in steady state exponential growth and 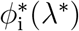 are the growth laws of eq. 31 32. The relation between *λ* and *σ* can be calculated from the ribosome growth law 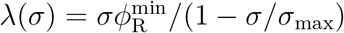 (see ref. [8] for details).

Here, we will employ the same approach in defining χ_H_ and 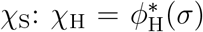 and χ_S_ = 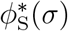. Therefore we need to know the parameters that specify the growth law for the survival and harmful sector, which are the slope and the offset. The issue is that these two sectors have not been identified yet and as any collection of proteins, they have their individual slope and offset, which are unknown. To avoid having to define slope and offset we eliminate the explicit dependence on the parameters. We can demonstrate that for any protein, or group of proteins (sub-sector), *i*, belonging to *ϕ*_up_ the sector-specific growth law described by eq. 31 can be obtained from the growth law of any other protein of the group, *j*, by applying a linear transformation to the first sector:

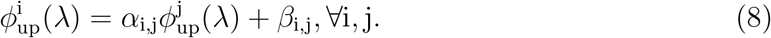

Therefore, we can use the fact that *ϕ*_H_ is a part of *ϕ*_up_ to write:

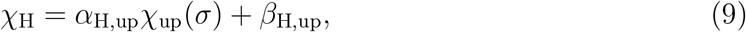

and the same logic applies to *ϕ*_S_, which is a part of *ϕ*_down_:

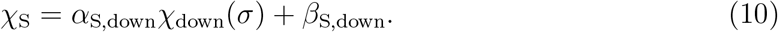

The definition of the regulatory function concludes the description of the global regulation model. For additional details on the definition and simulation refer to the methods section.

### Changes in harmful and survival sectors have identical impacts on adaptation if the sectors are globally regulated

In the above sections, we have defined the general structure of the model. In this section, we will further specify the adaptation dynamics of *ϕ*_H_ and *ϕ*_S_. According to the introduced theoretical framework, the dynamics of the size of the harmful and survival sector are determined by:

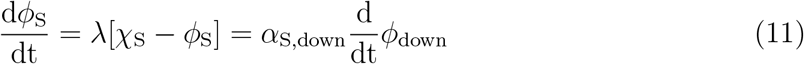

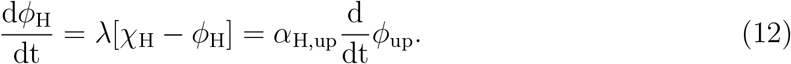

We can show that these two scenarios (production of survival proteins and dilution of harmful ones) are linearly related, and therefore have identical effects on death rate. This is a consequence of the constraints on the size of the sectors introduced in [3]:

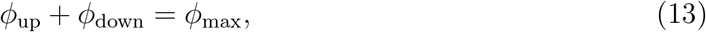

where *ϕ*_max_ is a constant, given by the size of the “housekeeping sector”, whose size does not change with the growth rate and stays constant in nutrient-limited growth. Because of how we define them, this constraint also holds for the regulatory functions, χ_down_ = *ϕ*_max_ − χ_up_, which allows us to derive 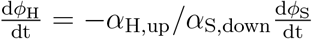, meaning that the dynamics of both sectors are inversely related. Equivalently, using the linear transformation (eq. 8), the proteome constraint (eq.13) and the relation between death rate and proteome sectors (eqs. 3 and 4) we can derive that both the “harmful” and the “survival” sector scenarios yield mathematically indistinguishable results

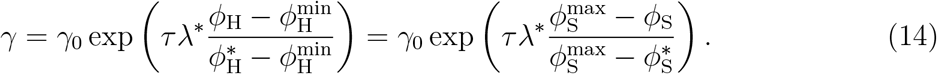

This means that if the adaptation follows the global proteome dynamics, it would be impossible to tell whether an adaptation in the death rate was caused by either a reduction of the harmful sector or an increase in the survival sector.

To better understand the global adaptation dynamics, fig. 3A-F shows an example simulation, chosen with parameters set of growth rate, time-point of nutrient running out, and steepness of nutrient depletion to highlight the dynamics. When the nutrient concentration (fig. 3A, orange line) decreases below the affinity of the uptake proteins, the growth rate decreases (fig. 3B, black line). In addition, as a result of the decreased uptake yet still high ribosome abundance, the translational activity *σ* decreases (fig. 3C, black line). Because in the FCR model the regulatory functions χ_up_ and χ_down_ are set by the translational activity, the regulatory functions adapt (fig. 3D, red and blue lines). Fig. 3D shows how they change with respect to their initial steady state value. To measure the adaptation of the proteome, we plot the relative proteome change concerning the steady state composi-tion 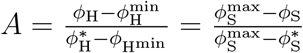, which determines the death rate (eq.14), in Fig. 3E. Because *A* = 0 would mean maximum adaptation, the predicted change from 1 to roughly 0.5 in the example simulation means that the bacteria can adapt about half of their total capacity. This is a consequence of the fact that sector size changes slower than regulatory functions.

**FIG. 3:**
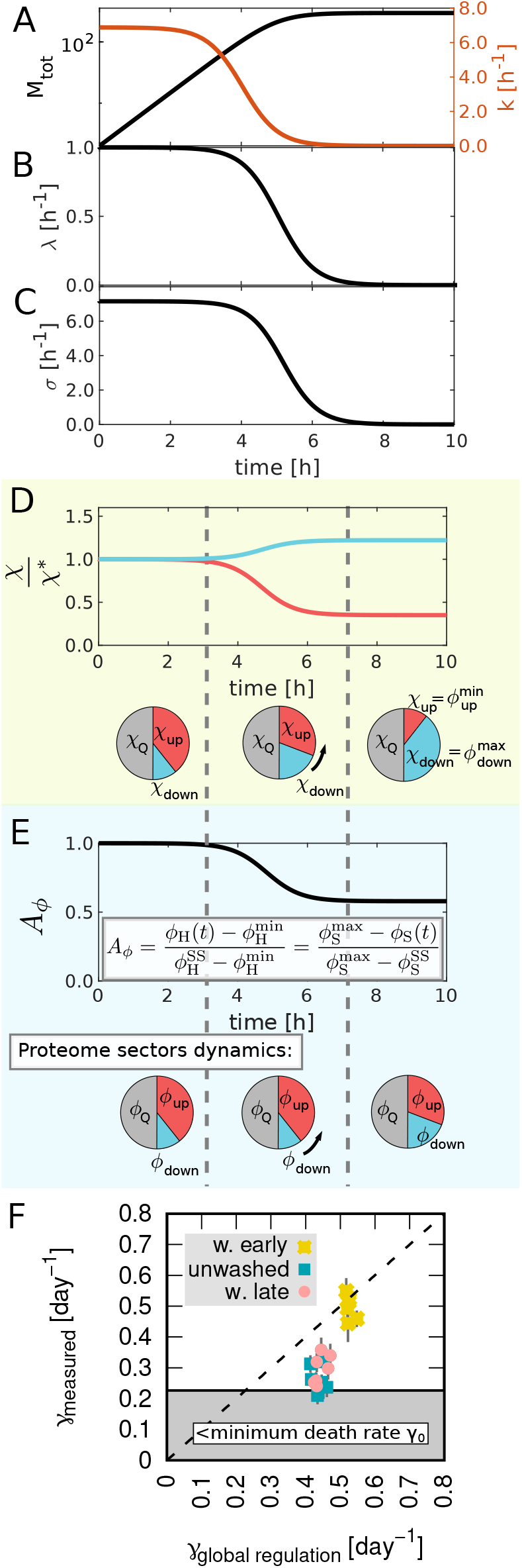
Global proteome regulation is not sufficient to reproduce the experimental death rate data. (A-E) Example of the global adaptation dynamics during entry into starvation. (A) Time courses of the mass of the bacteria culture *M* (black line) and nutrient affinity *k* (orange line) from steady-state exponential growth to stationary phase. The adaptation phase starts as the nutrient affinity starts to decrease and it stops when *k* reaches zero. (B) Time course of the growth rate *λ*, and (C) of the translational activity *σ*. Both quantities start to decrease as the nutrient affinity drops. (D) Time course of the regulatory functions of the up and down sectors *χ*_up_ (red bottom line) and *χ*_down_ (blue top line), normalized by their steady-state value. Beneath the plot, a graphical synopsis of the regulation functions dynamics, with arrows highlighting adaptation. The regulatory functions change following changes in *σ*. (E) Proteome adaptation extent (definition of the global adaptation in the inset). During the adaptation phase this term decreases and as a consequence, the death rate *γ*, which depends on it, decreases too. Beneath the plot, a graphical synopsis of the sector dynamics, with arrows highlighting adaptation. (F) Comparison of the experimental results with predictions. The points are 7 replicates of the experiment carried on using MM + Lactose 0.5 % as substrate. The y coordinate represents the measured death rate with uncertainty, while the x coordinate is the death rate predicted by our theory. The data points are for the washed early (yellow asterisks), washed late (pink rounds), and unwashed cultures (blueish squares). The dashed line is the bisector *γ*_measured_ = *γ*_global_, while the shaded area represents the region where the measured death rate is lower than the minimum possible one 0.21day^−1^ [2]. This plot confirms the exponential relation between growth and death rate for the washed culture but also shows that the global regulation hypothesis cannot explain the observed death rate of the two adapted cultures. In particular, the adapted cells are dying slower than expected, some of them approaching the minimum achievable death rate, meaning that their adaptation strategy is more efficient. Please refer to the Methods section for more details.

### Cell adapts faster than predicted by the global proteome regulation model

After having defined the theoretical model for the death rate, we turn to the comparison of the model predictions with our experimental observation. To simulate the adaptation dynamics of the experiments, we interpolate the data using a sigmoidal fit (fig.S 14), which we use as input for our simulation to obtain the composition of the adapted proteome. We recall that our model assumes that washing at any time during growth will freeze adaptation in the current state. As a final result of the simulation we obtain the predicted death rate, according to eq.14, and compare it to experimental data. Fig. 3F shows this comparison. In the case of a match between experimental data and simulation points should fall along the dashed diagonal line, defined by *γ*_data_ = *γ*_model_. The non-adapted (“washed early”) cultures (yellow asterisks) are scattered around the dashed line because they correspond to the sudden starvation of ref. [2]. On the other hand, the cultures that have adapted, “washed late” (pink circle) or “unwashed” (blueish squares), lay far away from the diagonal line, and show significantly lower death rates close to the minimum, *γ*_0_, obtained for prolonged growth (shaded area). This means that E. coli during entry to the stationary phase adapts more than expected from a simple, global proteome memory model.

One possibility to explain this phenomenon is that bacteria speed up the adaptation by specifically regulating the survival and the harmful sector, for example by transiently redirecting more/fewer resources towards *ϕ*_S_ and *ϕ*_H_, rather than regulating on the global level of up and down-sectors. We will explore this possibility in the next section using a “targeted” regulation model.

### Targeted up- and down-regulation of the survival and harmful sectors can improve cell adaptation

Because the global regulation model is not sufficient to reproduce the data, we introduce a new, “targeted” version of the regulation, that allows cells to direct more/less biosynthesis flux to *ϕ*_S_ and *ϕ*_H_ during the entry to stationary phase. To implement this, we add to the FCR model two parameters: *u*(*t*) ∈ (1, ∞) and *d*(*t*) ∈ [0, 1), which set how much *χ*_S_ and *χ*_H_ are up- or down-regulated in addition to the global regulation,

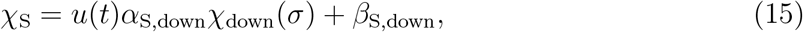

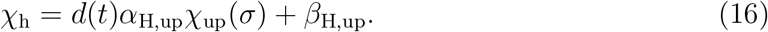

The two sectors are affected by the additional up- or down-regulation only during the entry to the stationary phase when cells sense the approaching starvation. When *ϕ*_S_ and *ϕ*_H_ reach their extreme values the additional up/down regulation is switched off. This means that under normal growth conditions *u*(*t*) = 1 = *d*(*t*), they acquire a greater or lower value during the transition, and when the S and H sectors have reached their target value they revert to 1. We call the values that *u*(*t*) and *d*(*t*) acquire during the transition *u*(*t*)^max^ and *d*(*t*)^min^. More specifically, these two parameters behave as:

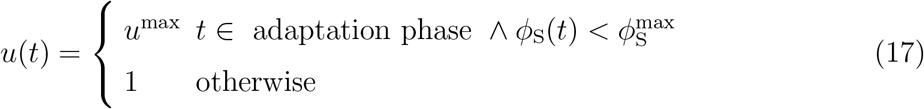

and

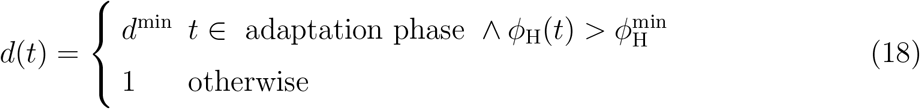

There are two consequences of this time dependence: (i) firstly the dynamics of *ϕ*_S_ and *ϕ*_H_ adapt quicker than the up and down sectors and (ii) because of the targeted regulation *ϕ*_S_ and *ϕ*_H_ are not anymore inversely related and thus their effect on death rate is not identical. This means that we can distinguish whether the determining sector is the “harmful” or “beneficial” one by checking which one of the two scenarios can describe the data,

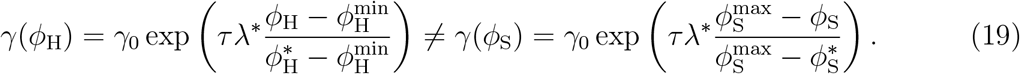

The overall adaptation dynamic is sketched in fig. 4A-E for an example simulation of an up-regulated survival sector and in fig. 4G-M for a down-regulated harmful sector. As before, the adaptation phase starts when the nutrient affinity decreases (orange line in fig. 4A). As a consequence, both the translational activity *σ* and the growth rate *λ* will drop (fig. 4B and C), and the proteome regulation will change accordingly. Fig. 4D shows how the regulatory functions change relative to their initial steady state value. Because of the targeted upregulation, the regulatory function of the survival sector (χ_S_, dark blue line) increases more than the regulatory function of the down-sector. To measure the adaptation of the proteome, we plot the relative proteome adaptation of the survival sector with respect to the steady state composition 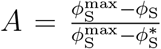, which determines the death rate (eq.19), in Fig. 3E. The plot shows that this targeted regulation of the survival sector (dark blue line) leads to more effective adaptation compared to an untargeted down-sector (black line). Therefore the death rate will decrease more with respect to the global regulation case, even if it does not reach complete adaptation (*A* = 0). Figure 4 panels G-M show the analog case for the harmful proteome sector *ϕ*_H_, which is down-regulated. Note that while the harmful sector is down-regulated to zero in Fig. 4L, the adaptation of the proteome fig. 4M (dark red) is considerably less than in the case of the targeted survival sector of 4E (dark blue). This is because the harmful proteome can only decrease by dilution, whereas the synthesis of beneficial proteome can in principle be up-regulated at will.

**FIG. 4:**
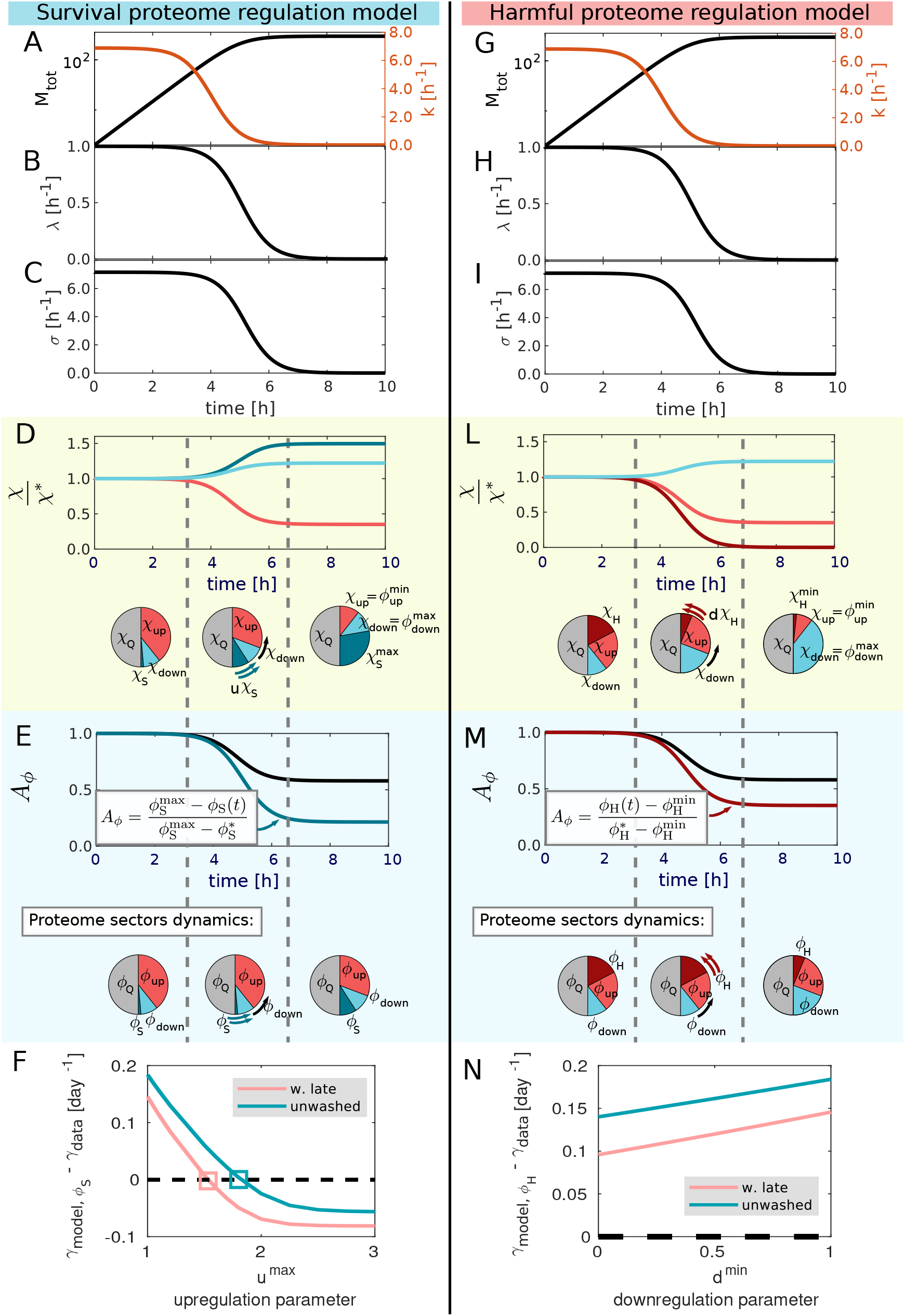
Targeted up-regulation of the “survival” proteome sector can explain the extent of adaptation to starvation displayed by the unwashed and late-washed cultures.(A-E) Example of the survival-targeted adaptation dynamics during entry into starvation. (A-C) Same as Fig. 3A-C: Dynamics of the mass *M*, nutrient affinity *k*, growth rate *λ* and translational activity *σ*. (D) Dynamics of the regulatory functions χ_up_ (red bottom line), χ_down_ (blue midline) and χ_S_ (dark blue top line), normalized by their steady-state value. Here χ_S_ follows the same dynamics as χ_down_ with the additional targeted up-regulation. (E) Proteome adaptation extent (definition for *ϕ*_S_ in the inset). This term (dark blue line) decreases faster than the adaptation of the other sectors (black line, same as in fig. 3F) because of the targeted regulation. Beneath D and E, a graphical synopsis of the shown dynamics. (F) Fit of the up-regulation parameter *u*^max^ (eq. 17). The plot shows the results of the targeted-regulation model, for different values of *u*^max^. The simulation parameters of the adapted cultures were extracted from the MM + lactose 0.05 % experiments. The y-axis shows the difference between the predicted death rate and the measured one, for different values of the x-coordinate *u*^max^ (*u*^max^ = 1 is identical the global regulation model). The plot shows that the up-regulation of *ϕ*_S_ is relatively low, with *u*^max^ between 1.5 and 2 (value of the intercept, empty squares). (G-M) Example of the harmful-targeted adaptation dynamics during entry into starvation. (G-I) Analogous to panels A-C. (L) Time course of the regulatory functions χ_up_ (red midline), χ_down_ (blue top line) and *χ*_H_ (dark red bottom line), normalized by their steady-state value. Here χ_H_ follows the same dynamics as χ_up_ with the additional targeted down-regulation. (M) Proteome adaptation extent (definition for *ϕ*_H_ in the inset). This term (dark red line) decreases faster than the adaptation of the other sectors (black line, same line as in fig. 3F) because of the targeted regulation. Beneath L and M, a graphical synopsis of the shown dynamics. (N) Fit of the value for the down-regulation parameter *d*^min^, eq. 18. The plot shows the results of the targeted-regulation model, for different values of *d*^min^. The simulation parameters of the adapted cultures were extracted from the MM + lactose 0.05 % experiments. The y-axis shows the difference between the predicted death rate and the measured one, for different values of the x-coordinate *d*^min^ (*d*^min^ = 1 is the global regulation model). The plot shows that this dynamics is not able to explain the data even in the extreme case of pure dilution (*d*^min^ = 0).

### Targeted up-regulation of the survival sector explains the observed death rates and adaptation dynamics

To understand which one of the two scenarios (up-regulation of survival proteins or down-regulation of harmful ones) can describe the adaptation during entry to the stationary phase, we compare the measured death rate and the one predicted by the two hypotheses for different values of the targeted regulation parameters *u*^max^ and *d*^min^. The results of this study are shown in Figure 4F and N. We plot the *u*^max^ and *d*^min^, the parameter of the “targeted” regulation models against the difference between the death rate of the survival & harmful proteome hypotheses and the average death rate of adapted lactose cultures. We find that for the survival proteome hypothesis, there exists a value of the up-regulation parameter *u*^max^, around 1.5 and 2, for which the death rate of the model equals the experimental data. On the other hand, the harmful proteome hypothesis is not able to reproduce the data on the adaptation extent for any value of the down-regulation parameter *d*^min^. In this scenario, the expected death rate is always higher than the measured one, even in the extreme case of pure dilution of all the harmful proteins, *d*^min^ = 0, because harmful proteome cannot diluted in the absence of growth (if we neglect degradation). These results show that of the two possible adaptation strategies, the only one that is efficient enough to reproduce the experimental data is the survival proteome up-regulation. The adaptation cannot rely simply on the dilution of harmful proteins, as this process is too slow to account for the observed death rate.

In addition, it is surprising that, according to the model, bacteria up-regulate their survival sector by just two-fold, given that individual operons can be up-regulated 100-fold [7, 32, 33]. From our data, we know that bacteria are not able to fully adapt, because death rates after adaptation are still higher than the minimum, Fig. 3F. Full adaptation could be achieved by up-regulating the survival sector more than 2-fold, or by degrading the harmful sector until is it completely gone. This suggests that lowering the death rate requires major resource allocation towards survival proteins and likely exceeds the simple up-regulation of a few operons, and that degradation, if it is used at all, is not able to completely degrade all the unnecessary or harmful proteins.

## DISCUSSION AND CONCLUSION

We presented an experimental study of the adaptation dynamics of *E. coli* during shifts into the stationary phase induced by carbon starvation, focused on its consequences for the death rate. We developed a mathematical model based on the FCR framework (a model of growth response to nutrient shift introduced in ref [8]) that describes the observed behavior by connecting the observed death rate to the proteome sectors, in particular introducing two distinct proteome categories: harmful and survival. Our hypothesis is that the adaptation of these sectors determines the bacterial population death rate. In particular, our data is consistent with a model that includes an active and targeted allocation of resources during the transition to the stationary phase enhancing the production of survival proteins. Importantly, this is opposite to the global regulation, where the proteome is changed in response to the lowering of nutrient concentrations and, by chance, the adapted composition also enhances starvation survival. Interestingly, our simulations indicate that the predicted up-regulation factor for the survival sector is relatively low, raising the question of why bacteria do not choose to further up-regulate survival proteins and achieve the minimum possible death rate (*γ*_0_). This might be attributed to additional constraints or trade-offs that bacteria face during adaptation, for example the energy cost of making certain proteins. Moreover, the fact that bacteria cannot fully adapt rules out the possibility of the complete degradation of all the harmful proteins, which could be due to the limited targeted degradation in *E. coli*, or other constraints. Note that this does not imply that degradation is not in place at all, just that it is not sufficient, or that it is not working towards the achievement of the minimum observed death rate. This relatively low up-regulation could also be advantageous: if bacteria live in an environment that frequently oscillates between feast and famine, being able to enhance survival by changing just a part of the previous proteome composition helps to regrow quicker when nutrients become available again [2].

The results of our study emphasize the need for additional experimental investigations to test and further refine our hypotheses. In particular, a detailed analysis of proteome composition during shifts toward starvation will provide valuable insights into the specific changes that occur in the sectors. Moreover, understanding the extent to which the FCR framework accurately describes bacterial responses to nutrient depletion remains an important area of exploration. This model was not tested for starvation and shifts to very low growth, and it could be that some of its assumptions, for example that protein synthesis is completely halted when the cells stop growing, are not accurate in this setting. Another area of exploration regards single-cell behavior. A recent work [34] studied the death dynamics of isolated cells and confirmed that these cells present stochastic first-passage-time dynamics of cell death. At the single-cell level, it is also well documented that cells often choose bet-hedging [35, 36] strategies or switch to persistence phenotype [37] to ensure the survival of the colony in fluctuating environments. Both these strategies rely on bacteria switching their growth mode in a stochastic way, while in this work we instead propose a deterministic change in protein expression. However, given the bulk nature of our data, we cannot rule out the fact that this collective behavior is the result of stochastic protein expression at the single-cell level. Another recent work [38] studies the protein expression of starved bacteria. The authors follow single-cell starvation dynamics with a microfluidic device and measure protein expression by tracking the amount of produced fluorescent proteins. They found that cells can change their intracellular protein concentrations during the first few hours into starvation. Among the twenty tested promoters, there are almost none whose protein concentrations remain the same as in the exponential growth, as this either increases or decreases. These results are in line with our hypothesis of targeted regulation of specific proteins upon the onset of starvation. In our model, we hypothesize that proteins cannot be produced after all substrate has run out, but this hypothesis can be easily relaxed by introducing a delay between growth arrest and protein production arrest, as ref [38] suggests. The main difference between our work and this new study [38] is that in their experiments they do not find any qualitative difference between cells that experience a smooth or an abrupt entry into starvation. We think that this discrepancy is due to the very different experimental setups utilized (batch culture VS microfluidic) and the way gradual starvation is induced (actual depletion VS low concentration).

In conclusion, our model provides a possible framework for the observed adaptation dynamics of Escherichia coli during shifts toward the stationary phase. The interplay between proteome sectors, degradation processes, and additional cellular and energetic constraints is a rich area for future research. In this context, we refer to several recent studies that quantitatively explored starvation physiology. First of all, this work is the natural follow-up of a series of papers focused on starvation survival [2, 39], and it gives experimental and theoretical support to the hypothesis of proteome-determined death rate introduced in ref. [2]. Recently, other works adopted this adaptation hypothesis, for example ref. [24] connected the death rate under antibiotics treatment to the size of a “stress sector” that is induced even during growth if the cells grow in stressful environments. The fact that our hypothesis can be used to explain death dynamics under very different sources of stress (antibiotic treatment VS carbon starvation) proves its generality and its predicting power. More in general, in the last years the dynamics of the stationary phase have collected a lot of attention, either because this phase is heavily related to other important health-related questions, for example antibiotics survival [40, 41]. Another reason is that a phenomenological and quantitative understanding of what is happening during starvation and death is still missing, and in this context, other works shed light on how starving bacteria change in size, morphology, growth, and expression profiles [42] and the role of stochasticity in cell death [43]. Lastly, recent studies [23, 44] discovered a connection between cell wall robustness and death rates, and another study [45] shows that the proteins responsible for building the cell wall scale negatively with growth rate. These results imply that wall proteins belong to the “down-regulated” sector, the same as the survival proteins, and therefore they provide a potential candidate for it. Furthermore, another recent study [46] offers valuable insights into the relationship between membrane properties and bacterial proliferation. It could provide complementary information about how the expression of membrane proteins is influencing bacterial growth rates and, as a consequence of the *γ*-*λ* relation, also bacterial death rates.

## MATERIALS AND METHODS

### Data and code availability

The code used for the simulation of the models is available in an online repository: DOI: 10.17632/dz6hpkrzm3.3. The raw data are all shown in the supplementary table S1, and they are plotted in the supplementary figures.

### Experimental procedure

#### Strains

The strain used in this study is the wild-type *E. coli* K-12 strain NCM3722 [47].

#### Growth medium

The minimal medium used in this study is based on three ingredients [48]: (i) N-C-Mg-medium containing K_2_SO_4_ (4g), K_2_HPO_4_ (54g), NaCl (10g), KH_2_PO_4_ (18.8g) in one liter of deionized water (ii) NH_4_Cl 1M and (iii) MgSO_4_ *·* 7H_2_O 4%. To make 45 ml of minimal medium we mix 32.65 ml of deionized water, 11.25 ml of N-C-Mg-medium, 900 *µl* of NH_4_Cl 1M and 112.5 *µl* of MgSO_4_ *·* 7H_2_O 4%. The nutrient sources (i.e. lactose, CAA etc.) were added to this minimal medium in the prescribed concentrations in each experiment.e

#### Growth and washing protocol

For growth experiments, the same procedure described in ref. [39] was applied. We recall here the main steps. Before each experiment, we took the cells from the -80^°^C stock, placed them on an LB agar plate, and grew them at 37^°^C for one day before putting the plate in the 4^°^C fridge. Growth was then carried out in three steps: seed culture, overnight culture, and experimental culture, each cultured at 37^°^C in a water bath shaker at 250 rpm with water bath preservative. The goal of the seed culture is to grow the cells taken from the colony on the stock plate. The seed cultures are prepared with fresh LB medium and inoculated with a single colony from the stock plate. For every experimental condition, we prepared at least two different replicates. After 3-4h of growth, a sub-sample of the seed culture is washed via centrifugation (at 7500 rpm for one minute), suspended in the minimal medium, and inoculated in a new tube for the overnight culture. The size of the inoculum is calculated in order to have the cells still growing exponentially on the morning of the experiment. For this step, the medium used is the same as the third step. Cells perform at least ten doublings in the overnight culture. For the third step, a sample of the overnight culture is inoculated in the tube for the experimental culture, the amount chosen such that after three doublings the optical density of the inoculated culture is roughly equal to 0.05. After the three doublings we started measuring growth: at each time point a 200 *µ*l sample was extracted and optical density was measured using a 10 mm path length Sub-Micro Cuvette (Starna, Ilford, United Kingdom) at 600 nm in a Spectrophotometer (Genesys 20, ThermoScientific. Waltham, MA, USA). Measurements were taken every half hour for the exponential growth, and every 15 minutes during the transition phase. For the overnight culture, we grow 5 ml of culture in 20 mm x 150 mm glass test tubes (Fisher Scientific, Hampton, NH, USA) with disposable, polypropylene Kim-Kap closures (Kimble Chase, Vineland, NJ, USA). For the experimental culture, we grow 9 ml of culture in 25 mm x 150 mm glass test tubes. The washing steps are performed during the growth experiment. In particular, during the exponential phase for all the “washed early” (non-adapted) cultures and right after the transition phase for the “washed late” (adapted) ones. To wash the culture, all the culture in the experimental tube is transferred to a plastic tube, which is centrifugated at 3000 rpm at 37^°^C for 10 minutes. After the centrifugation step, the supernatant is removed and substituted with minimal medium, without nutrients. The culture is transferred in a new experimental tube and placed back in the heat bath.

#### Starvation protocol

For starvation experiments, the same procedure described in ref. [39] was applied. We recall here the main steps. After the cultures (adapted or non-adapted) reached the stationary phase, we waited for a time roughly equal to two doubling times before starting the starvation measurements. This was to let the cells complete the last division before starting the viability measurements. Viability was measured by plating on LB agar and counting the colony-forming units (CFU) after an incubation period of 24 hours at 37 degrees. Samples were diluted in fresh minimal medium without carbon substrate and spread on three LB agar plate dishes (92 × 16 mm, Sarstedt, Nuembrecht, Germany) using Rattler Plating Beads (Zymo Research, Irvine, CA, USA). The dilution step aims to have between 100 and 200 CFU for each dish. LB agar was supplemented with 25 mg=ml 2,3,5-triphenyl tetrazolium chloride to stain colonies and increase contrast for automated colony counting (Scan 1200, Interscience, Saint-Nom-la-Bretèche, France). Staining or automation of counting had no significant effect on viability measurements or accuracy, compared to un-stained, manually counted samples (*<* 1% systematic error). The plating-counting steps were performed once a day for approximately ten days, in order to avoid the emergence of mutations [49]. The value of the death rate *γ* is obtained by fitting the viability data points: viability(*t*) = *A* exp(−*γt*), with A and *γ* the parameters of the fit.

### Simulation of the experiments

To predict the adaptation of the sectors during the entry into the stationary phase, we simulate with the FCR model the observed experimental dynamics, using a custom-written MATLAB script that follows this procedure (link). The core equations which compose the model are the following:

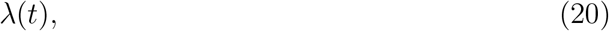

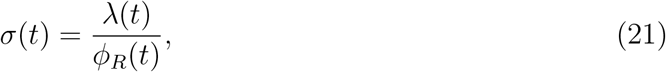

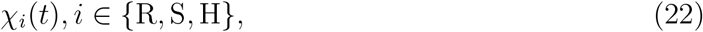

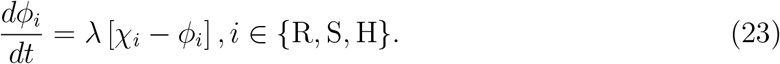

The equation *λ*(*t*) is the experimental input of the simulation. For each experiment, we fitted the time course of the growth rate with a sigmoidal function:

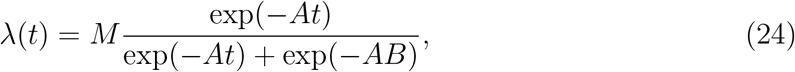

where the parameters M, A, and B represent the steady-state growth rate, the steepness of the transition to the stationary phase and the time at which the transition occurs respectively.

The dynamics of all the sectors can be derived from the ribosomal sector dynamic, as explained in the following method section, where we also introduce the functional form of the regulatory functions.

### Relations between the sectors

In this section, we will explain how to derive the dynamics of all sectors from the ribosomal one, and how to get rid of the unknown parameters *α*_i,j_.

Because of the linear relation that holds between all the sectors belonging to *ϕ*_up_ (and *ϕ*_down_) we can use the fact that both the ribosomal and the harmful sector are two subsets of *ϕ*_up_ to define the regulatory function χ_H_,

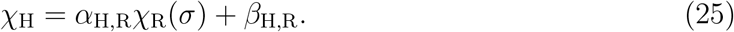

This has the advantage that χ_R_ is known from a previous work [8]: 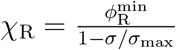, therefore the only unknown parameters left are *α*_H,R_ and *β*_H,R_. The same logic can be applied to *ϕ*_S_ and *ϕ*_P_, which are both part of *ϕ*_down_:

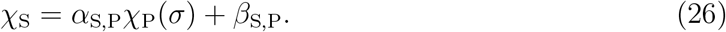

By using the constraint on the sector sizes introduced in ref. [3]: *ϕ*_R_ + *ϕ*_P_ = *ϕ*_*max*_ ≃ 0.55, we define χ_P_ = 0.55 − χ_R_. We have obtained that the regulatory functions of the survival and harmful sector can be derived from the one of the ribosomal sector, we now show how the values of *α*_H,R_, *β*_H,R_, *α*_S,P_ and *β*_S,P_ are irrelevant. By looking at eq. 12 and eq. 11 we can see that the value of the two *β* is not important because it cancels out when we take the derivative. To get rid of *α* we can define 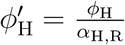. This sector evolves in the following way:

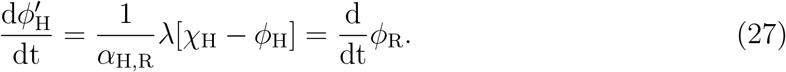

Therefore in order to know the dynamics of 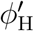 we do not need to know the value of *α*_H,R_, and if we use this sector 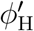 to calculate the death rate we will obtain the same result we would have obtained by using *ϕ*_H_:

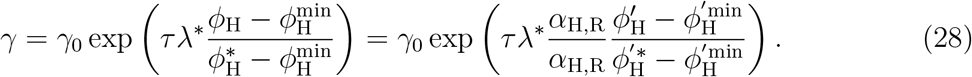

The same procedure can be applied to *ϕ*_S_ in order to get rid of *α*_*S,P*_.

Lastly, we define the two parameters *u*(*t*) and *d*(*t*), needed for the simulation of the targeted regulation model. Because it is not possible for MATLAB to integrate a stepwise function, for the two parameters *u*(*t*) and *d*(*t*) we define the following functions:

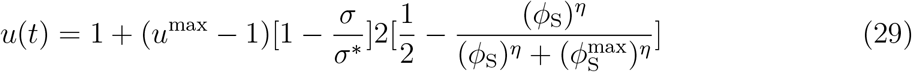

The first term in the squared parenthesis, 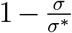 is equal to zero during steady state growth, and it ensures that *u*(*t*) part from 1 only during the transition phase. The term in the second parenthesis, 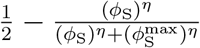 shuts down the up-regulation when 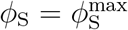, with *η* setting the sharpness of the transition (value used in the simulations *η* = 75). The parameter *d*(*t*) has a similar definition:

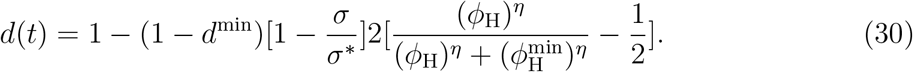

Supplementary figure S15 shows the trend of *u* and *d* versus *σ* and *ϕ*.

### Derivation of the “proteome memory model” from the growth laws for the up- and down-sectors

We hypothesize that the up- and down-regulated sectors depend linearly on the growth rate, like all the other defined sectors [4]. Therefore, the relation between growth rate and sector size can be expressed as

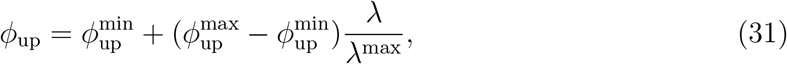

and

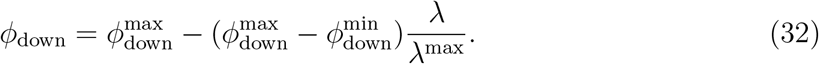

In both equations, *ϕ*^max^ and *ϕ*^min^ are the maximum and minimum size of the sectors, which are sector-specific, while *λ*^max^ is the theoretical maximum growth rate [3]. This functional form is given by the formula of a straight line passing through two points: (*λ* = 0, 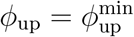) and (*λ* = *λ*^max^, 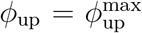) for eq. 31, and (*λ* = 0, 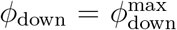) and (*λ* = *λ*^max^,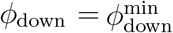) for eq. 32. These growth laws pass through all of the possible steady-state configurations: (*λ*^*^, 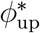) and (*λ*^*^,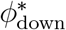), therefore if we choose a specific steady state point we can rewrite them in the following way

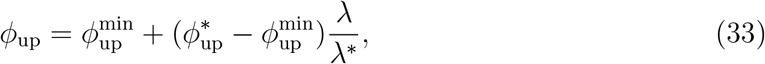

and

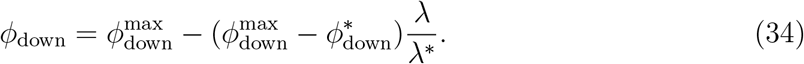

This re-definition is particularly advantageous for our scopes, because by inverting them we can obtain an expression for *λ*(*ϕ*_up_) and *λ*(*ϕ*_down_) that contains the adaptation extent:

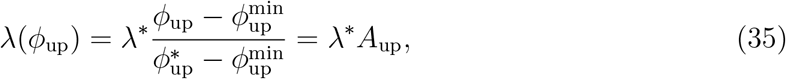

and

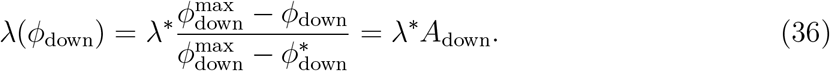

By inserting these two expressions in eq. 1 we obtain the protein memory model described by eq.s 3 and 4.

### Demonstration of the linear relation between the sectors

The up and the down sectors are composed of different subsets (for example *ϕ*_H_ and *ϕ*_R_ in *ϕ*_up_) which are regulated in a sector-specific way (i.e. their growth law is characterized by sector-specific slope and offset). It is possible to demonstrate that, if the expression of such subset is structured in a way that those sectors reach their extreme values when *λ* is either 0 or *λ*^max^, then exist a linear transformation between the growth laws. Indeed, two growth laws can be written as

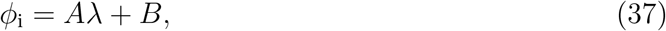

and

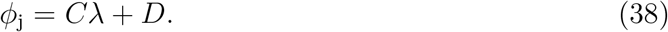

We now seek for a linear function of *ϕ*_i_, let us call it *f* (*ϕ*_i_), such that *ϕ*_j_(*λ*) = *f* (*ϕ*_i_(*λ*)). Given that *f* is linear, we can write it in this way: *f* (*ϕ*_i_) = *ηϕ*_i_ + ζ. If we take *η* = *C/A* and ζ = *D* − *B · C/A*, then the request *ϕ*_j_(*λ*) = *f* (*ϕ*_i_(*λ*)) is fulfilled.

An important exception of this demonstration is the catabolic sector *ϕ*_Cat_, which is a “down” sector that reaches it extreme values in the points (*λ* = 0, 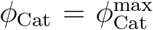) and (*λ* = *λ*^C^,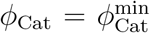), with *λ*^C^ = 1.17h^−1^, a value much lower than *λ*^max^ [28]. Because of this, for this sector, the linear relation with all the others does not apply.

## AUTHOR CONTRIBUTIONS

UG conceived the study. All authors designed the experiments. RD conducted the experiments with assistance from ZG, HA. RD and SJS designed the models. RD analysed the data and performed the model simulations. UG, SJS, and RD wrote the paper.

## ACKNOWLEDGMENTS

We are grateful to the members of the Gerland lab for useful discussions. We thank Marco Cosentino Lagomarsino and the members of his group for feedback on this work. We thank Gabriele Micali, Luca Ciandrini and Theo Gervais for feedback on the manuscript.

## SUPPLEMENTARY TEXT

*Dependence of the death rate on the substrate features*. Fig. S1 shows an important theoretical result, which we have used to understand the outcome of the experiments performed with substrates different from lactose. The plot shows the dependence of the difference in death rates of the adapted and non-adapted cultures, assuming that the adaptation follows the global regulation model. Therefore the result shown is more conservative than the result expected for real experiments, where we expect the survival proteome to be up-regulated. The difference in death rates is a crucial quantity that determines the experiment’s success. Indeed, if one experiment shows no difference between the starvation survival of the adapted and non-adapted cultures we cannot use it to test our hypothesis. This happens because the entry into the stationary phase is not long or compelling enough for the cells to adapt. To understand how this depends on the substrate features, we set up a simulation of cells growing on a single substrate, which they deplete. The substrate is characterized by *λ*_max_, which is the maximum growth rate achievable on the substrate, and *K*_half_, which is related to the substrate affinity. These two parameters determine the time course of the growth rate as the substrate *S*(*t*) gets depleted:

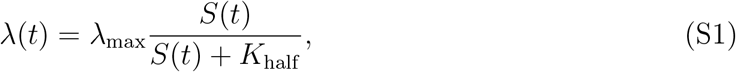

therefore they play a role in the dynamics of the entry into the stationary phase. As fig. S1 shows, the higher difference in death rates is achieved for substrate characterized by a high growth rate and a high *K*_half_ (so low affinity) like the lactose.

This result helps us understand why some of the performed experiments (in particular the one done with glycerol and acetate, fig.s S3 and S11) were not successful.

*Experimental data* Here we report all the data collected for this project, with a case-by-case commentary on why some data was included in the analysis and some was not.

The experiments performed with lactose were the ones that gave us the best results, and this is why in the main text we mainly refer to them. The results for the growth and starvation experiments are shown in fig. S2. The figure shows that the death rate of the adapted cultures (pink rounds and blueish squares, in the right column) is much less than the one which cannot adapt (yellow asterisks), as expected by our hypothesis. As shown in the main text, the adaptation extent is explained by the targeted up-regulation of the survival sector *ϕ*_S_.

We have found positive results also using other types of substrates, in particular with glucose, glycerol, casaminoacids (CAA), and lysogenic broth (LB). The results of the growth and starvation experiments carried out with these substrates are shown in fig. S3 - S6. In all of these experiments, we have observed a difference between the death rate of adapted and non-adapted cultures, like in the lactose case. We have carried out the analysis of these data with both the targeted and global model, to verify if the result holding for lactose (i.e. the confirmation of the survival sector hypothesis) is confirmed also by these other experiments. The experiments with glucose confirm our hypothesis, as fig. S7 shows. Also in this case, the global regulation model cannot explain the death rate of the adapted cultures, which is close to the minimum achievable value (fig. S7A). The analysis with the two targeted models (for the survival and harmful sector) shows that also in this case the only hypothesis that can explain the data is the survival sector one (fig. S7B), as the down-regulation of the harmful proteome is not enough (fig. S7C).

In the experiments with glycerol even if the difference in the observed death rate is measurable it is not enough to discriminate between the two hypotheses, as fig. S7A-C shows. Indeed here the difference in death rates is less pronounced than in the lactose and glucose case (fig. S3). This is a consequence of the lower growth rate achievable on glycerol, for two reasons: (I) the proteome composition associated with a lower exponential growth rate is similar to the one optimized for survival (indeed the survival sector belongs to the “down” sectors), (II) a lower growth rate imposes a cap on the adaptation extent, as eq. 10 of the main text shows. This is also confirmed by fig. S2, which shows that for low growth rates, we expect a small difference between death rates.

The experiments with CAA seem to confirm our hypothesis, as fig. S8 shows. Indeed in this case we see that the targeted up-regulation of the survival sector is the only hypothesis that can explain some of the data (fig. S8B and E, while the harmful sector one cannot. We can explain just part of the results because some of the cultures of this experiment display a death rate lower than *γ*_0_, which should be the minimum possible value in these growth conditions [2]). These data points lay outside of the range of our theory, therefore neither of the models can explain them. Given that CAA is a rich medium, we suppose that in this case, the cell can adapt in a way that is not captured by our theory. The FCR model, which we used to model the change in proteome composition during the entry into the stationary phase, has not been built to explain growth on rich medium, but only on carbon-limited substrates.

We have found a similar situation for the experiments with LB (fig. S9). Also in this case, the survival sector hypothesis is able to explain some of the results, but some cultures display a death rate that is too low to be captured by our model. Moreover, for this substrate, we were not even able to recapitulate the steady-state results. Indeed, if we look at fig. S9A-D-G, we can see that the yellow star-shaped points do not lay on the dashed line, meaning that the washed early cultures do not follow the steady-state relation between death and growth rate. This is possibly due to the fact that we did not achieve steady state exponential growth in LB, as shown by the mid column of fig. S6.

The last two experiments we performed, using acetate and mannose as substrate, are shown in fig. S10 and fig. S12 respectively. We did not carry out the numerical analysis on them, because they were not informative as the other experiments. Indeed, in both these experiments the adapted and non-adapted cultures display nearly identical death rates. As in the glycerol case, this is a consequence of the low growth rate achievable on these two substrates, which limits the adaptation.

### Details on dilution experiment

We designed this experiment to create a situation, where a bacterial culture grows rapidly and then displays a slow transition to stationary phase, starting from the entry down a high initial growth rate and a long entry into the by changing the external environment. These two features combined will allow the cells to better adapt their proteome, as fig. S1 shows.

Our strategy relies on two pieces of experimental evidence, shown in Fig. S13:

I. The relation between growth rate and mannose concentration is not constant for high concentrations. Instead, we found that the growth rate keeps increasing as the concentration increases (fig. S13B). Therefore, in this case, starting from a higher concentration will increase the starting growth rate;
II. The length of the transition phase depends on the initial concentration. The curves shown in Fig. S13B represent different concentrations thus their starting points are different and they are given by the highest concentration (the starting one). Thus the curve is longer the higher the concentration. Fig. S13C-F show that the time it takes for a culture to consume all the substrate is roughly the same for all concentrations. As a consequence, the culture that starts growing on a lower concentration spends a longer time when the growth rate is decreasing, which is the entry into the stationary phase.

We want to design a way to combine the higher growth rate of high-concentration environments with the long entry into the stationary phase of the low-concentration ones. The way to jump from the high-concentration environment to the lower-concentration one is the dilution of the experimental sample, where each dilution step lowers both the optical density and the substrate concentration. We performed the experiment by using the following concentrations: 0.200%, 0.100%, 0.050%, 0.025% and 0.010%. We started growing the cells on minimal medium + mannose 0.200%, and we diluted the sample each time the OD_600_ reached a value of 0.2 (before the growth started to slow down). We follow the growth dynamics by measuring optical density at regular time steps. We washed part of the culture at the time of the first dilution. After having performed all the dilution steps we obtain a culture growing in mannose 0.010%, which is left to consume all the substrate and enter the stationary phase. Then we followed the starvation dynamics of the washed and unwashed culture.

## SUPPLEMENTARY TABLE

**TABLE S1:**
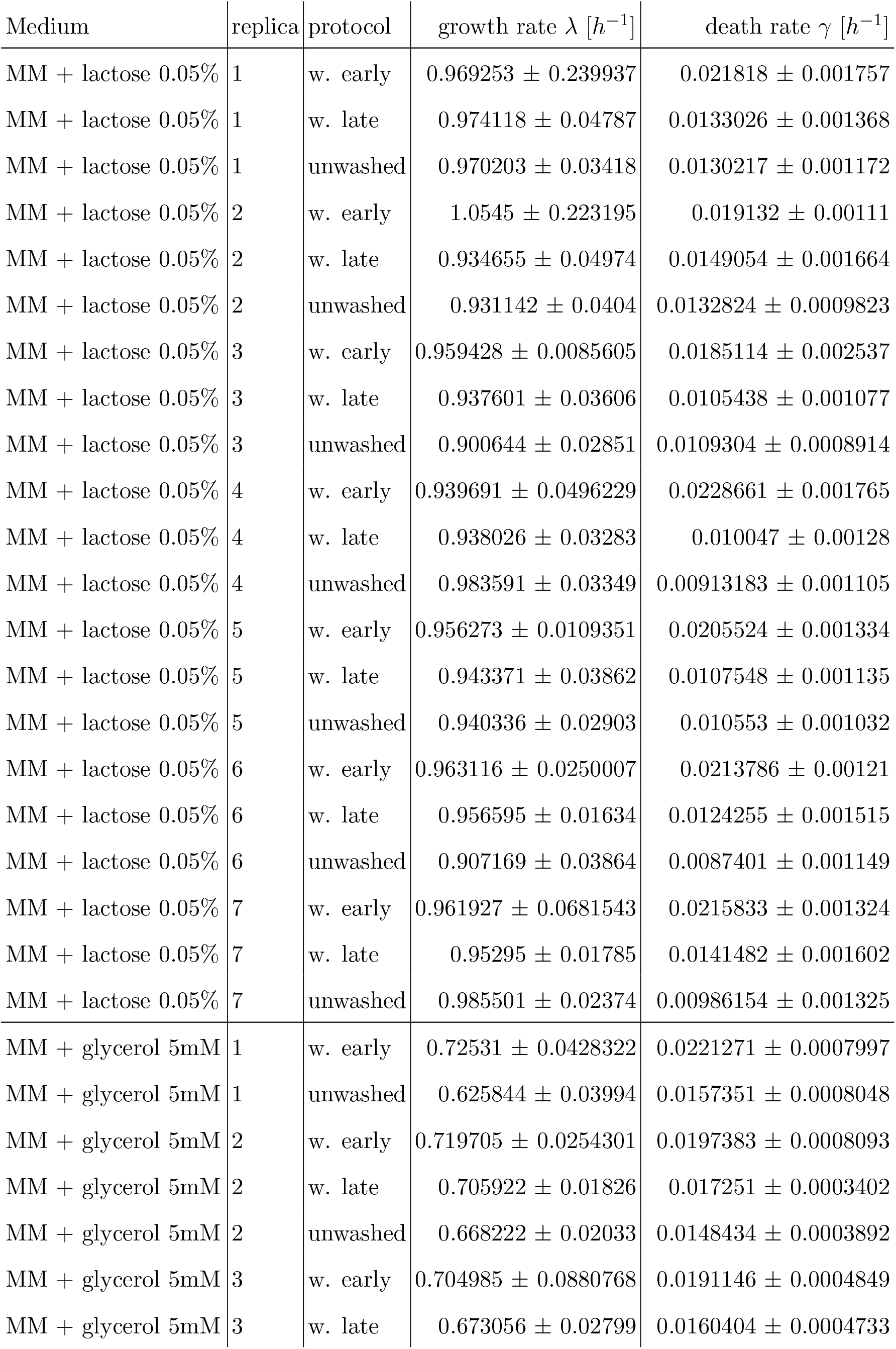
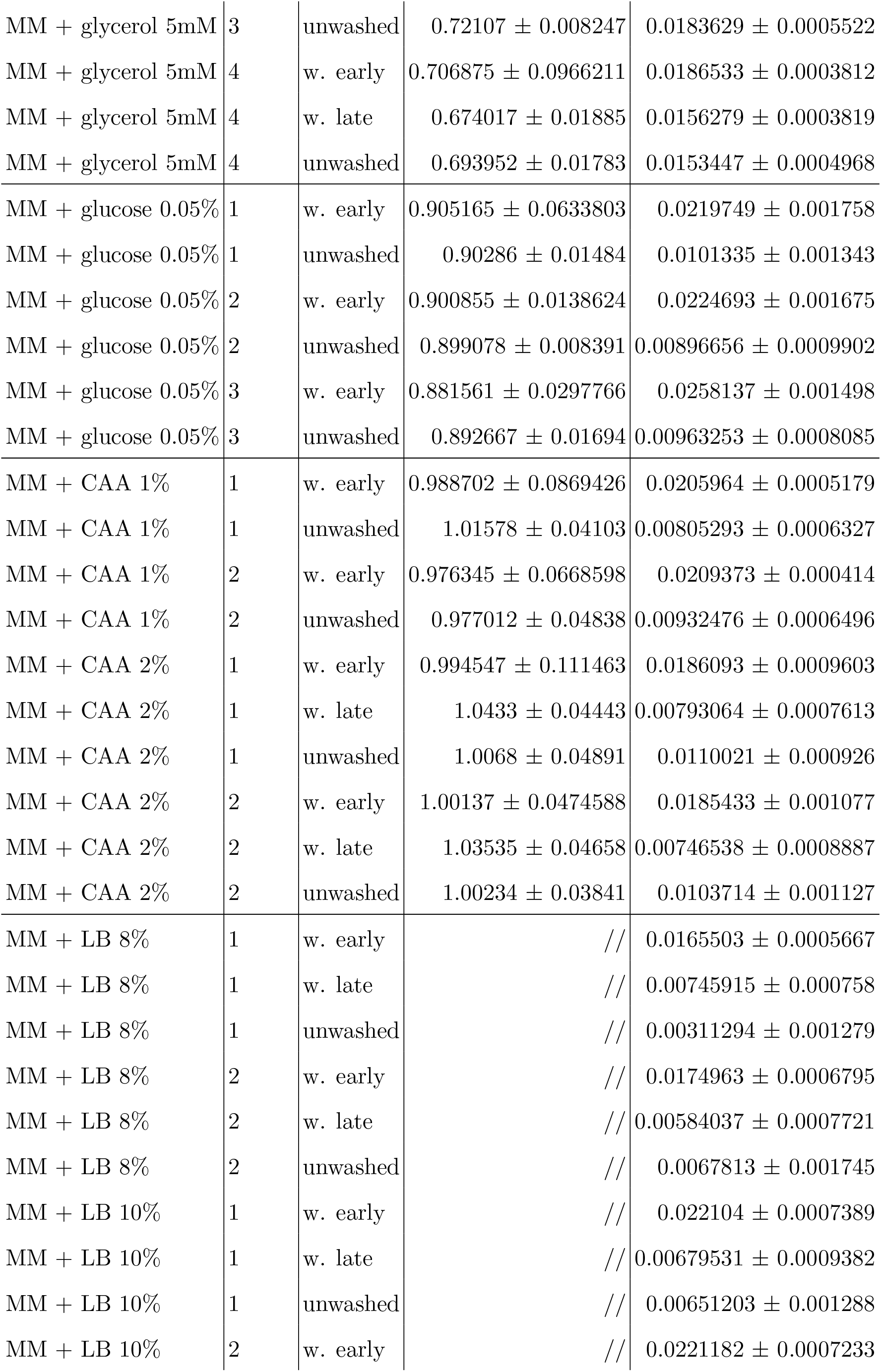
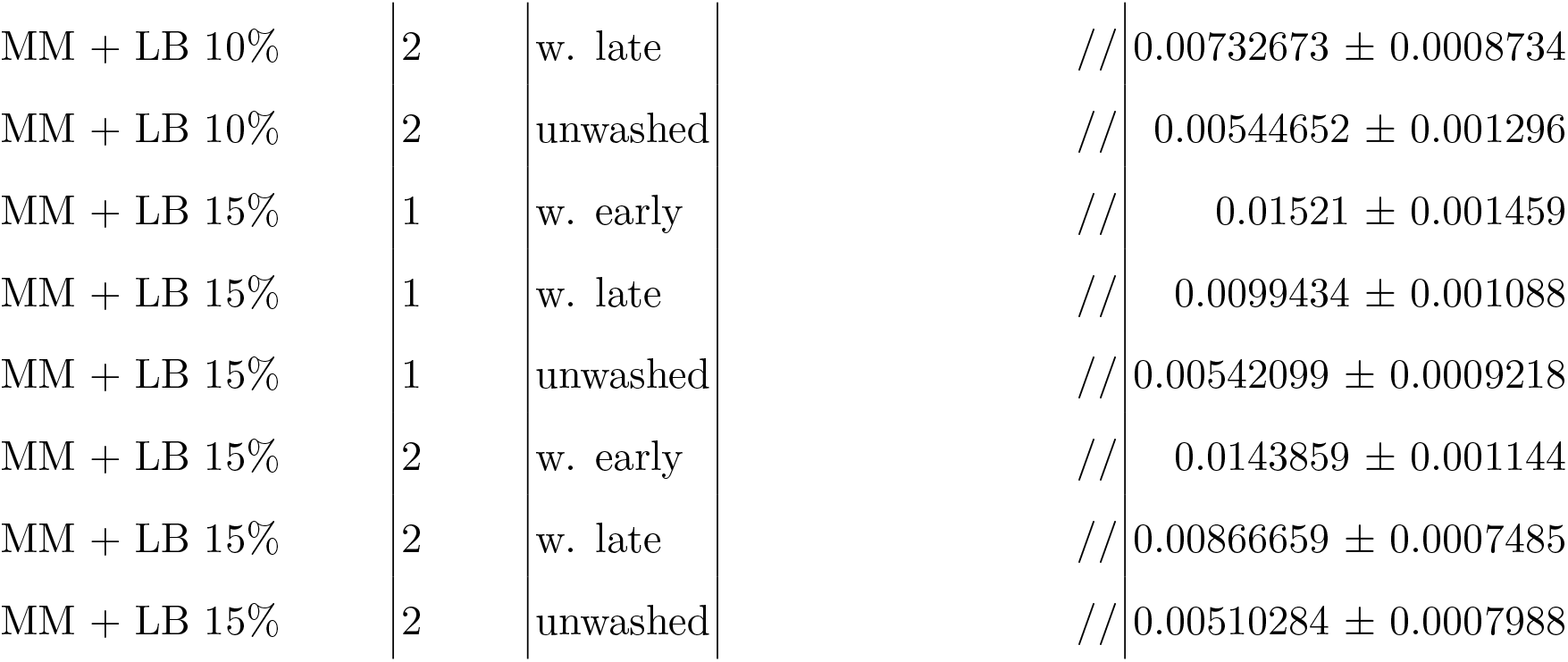
The collected experimental data for growth and death rates in various media for the three types of cultures: non-adapted (washed early) and adapted (washed late and unwashed). The growth rate for the non-adapted cultures is given by the average of the observed values, displayed with the absolute error. The growth rate of the adapted cultures is given by the fit of the sigmoidal function of eq.24, displayed with the fit uncertainty. The death rate of all cultures is given by a linear fit of the log-transformed data, displayed with the fit uncertainty. The growth rate value is missing for the experiments conducted in MM +LB at every concentration because cells do not display steady-state exponential growth in that case (shown in fig.S6)

## SUPPLEMENTARY FIGURES

**FIG. S1:**
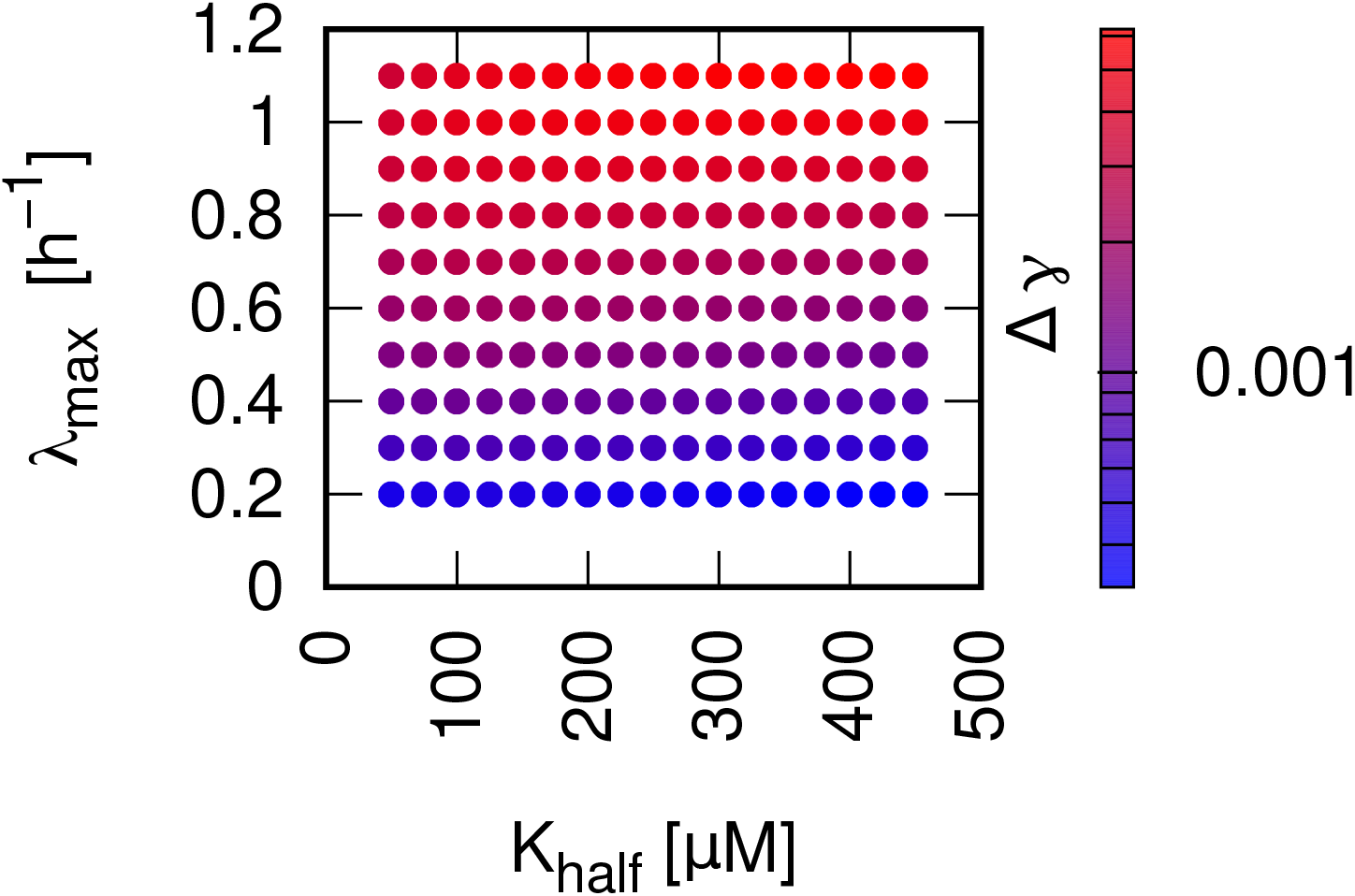
Dependence of the death rate on the substrate features. The figure shows the results of the numerical simulation of the global regulation model for an environment characterized by a single substrate running, which runs out as cells feed on it. The features of the substrate are the maximum growth rate the substrate supports, *λ*_*max*_, and the substrate affinity *K*_*half*_. Each point represents a single simulation, the x-coordinate shows the value of *λ*_*max*_ for that simulation, while the y-coordinate the value of *K*_*half*_. The color code shows the difference in death rates *γ*^*^ −*γ*^*adapt*^, where *γ*^*^ = *γ*_0_ exp(*τλ*^*^) and 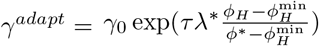. The plot shows that the difference in death rates between the adapted and the non-adapted culture depends on the features of the substrate. In particular, the higher are *λ*_*max*_ and *K*_*half*_ the wider the gap in the two death rates.

**FIG. S2:**
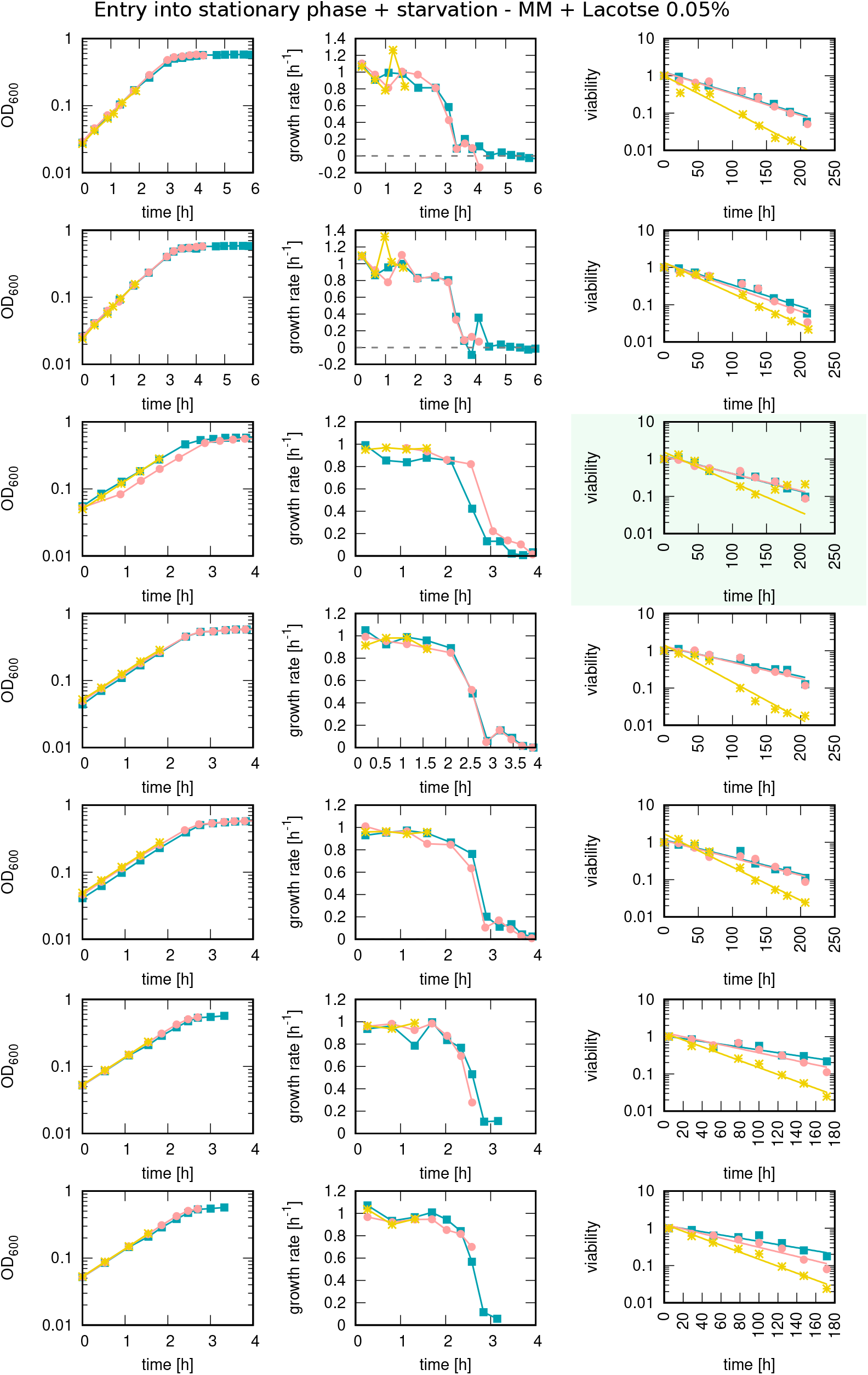
Measurements of OD_600_, growth rate, and starvation survival for the 7 independent replicates of the experiment with minimal medium + lactose 0.05%. Each row represents an independent replicate of the experiment, with three cultures for each replicate: “washed early” (yellow asterisks) “washed late” (pink circles), and “unwashed” (blue squares). For each row, the first plot on the left shows the time course of the mass of the colony, measured via the OD_600_, which is shown on the y-axis. These plots show that for all three cultures, the mass of the colony grows exponentially, slows down during the entry into the stationary phase, and then stops growing. The plot in the middle shows the time course of the instantaneous growth rate, with the growth rate on the y-axis and the time on the x-one. The growth rate starts constant during steady-state exponential growth and it drops to zero during the entry into stationary phase. The growth rate is computed from the OD_600_ data in the following way: 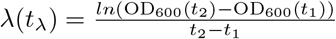, with 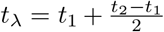. The plot on the right shows the survival dynamics of the starved cultures. The points represent viability measurements, the y coordinate shows the viability with respect to the first day of starvation, in percentage, and the x coordinates the time into starvation at which the measurement is performed. The solid lines are exponential fits of the data points. The viability plots show that the death rate of the washed early cultures (those who cannot adapt) is higher than the one of the other two for all the replicates of the experiment, as expected. Note that in one of the experiments, highlighted in green, after roughly 150 h into starvation, we notice the emergence of a mutant in the washed early culture, which increases in an anomalous way the viability measurements (yellow asterisks). For this reason, for that experiment, we performed the fit for the death rate only using the points before the emergence of the mutation.

**FIG. S3:**
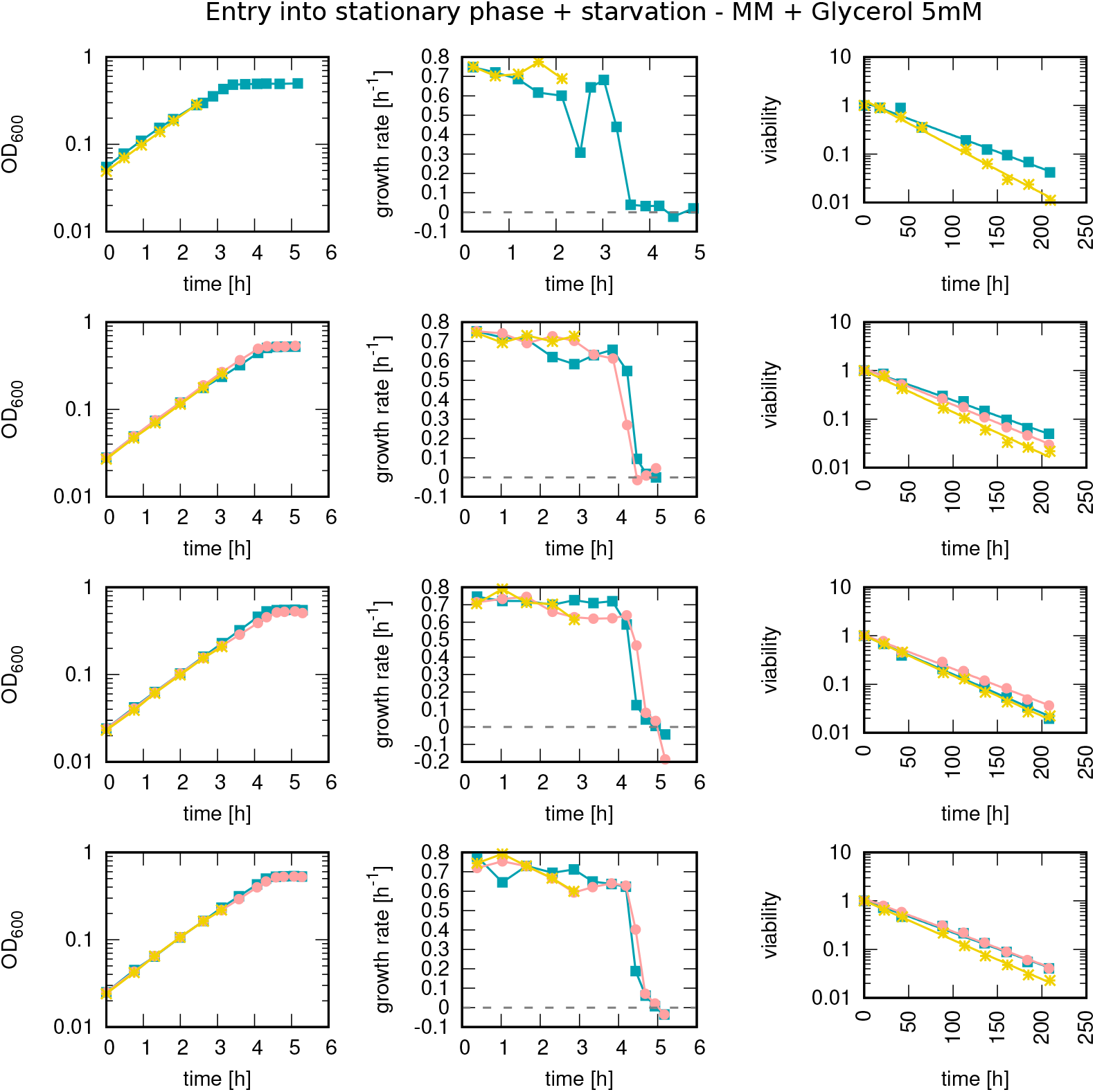
Measurements of OD_600_, growth rate and starvation survival for the 4 independent replicates of the experiment with minimal medium + glycerol 5 mM. Like in fig. S2, each row represents an independent replicate of the experiment with the three cultures: “washed early” (yellow asterisks) “washed late” (pink circles), and “unwashed” (blue squares). For each row, the first plot on the left shows the time course of the mass of the colony, measured via the OD_600_, which is shown on the y-axis. The plot in the middle shows the time course of the instantaneous growth rate, with the growth rate on the y-axis and the time on the x-one. The growth rate starts constant during steady-state exponential growth and it drops to zero during the entry into the stationary phase. The growth rate is computed in the same way as for Fig S2. The plot on the right shows the survival dynamics of the starved cultures. The viability plots show that the death rate of the washed early cultures (those who cannot adapt) is slightly higher that the one of the other two for all the replicates of the experiment. The first replicate does not have the “washed late” culture because at that stage we did not include the second washing step in the procedure yet.

**FIG. S4:**
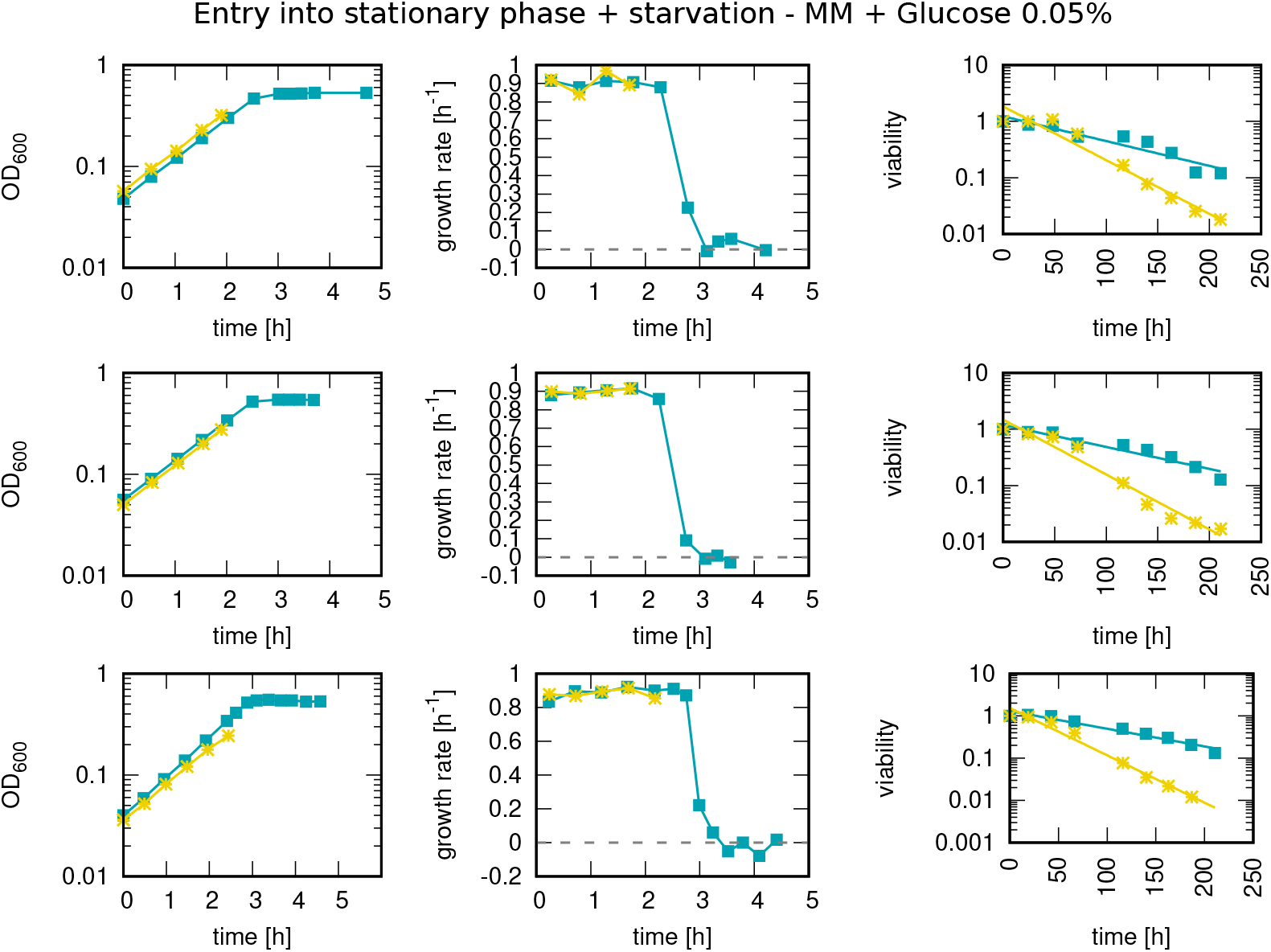
Measurements of OD_600_, growth rate, and starvation survival for the 3 independent replicates of the experiment with minimal medium + glucose 0.05%. Each row represents an independent replicate of the experiment with “washed early” (yellow asterisks) and “unwashed” cultures (blue squares). For this substrate, we do not have the “washed late” culture because at that stage we did not include the second washing step in the procedure yet. For each row, the first plot on the left shows the time course of the mass of the colony, measured via the OD_600_, which is shown on the y-axis. The plot in the middle shows the time course of the instantaneous growth rate, with the growth rate on the y-axis and the time on the x-one. The growth rate starts constant during steady-state exponential growth and it drops to zero during the entry into the stationary phase. The growth rate is computed in the same way as in fig S2. The plot on the right shows the survival dynamics of the starved cultures. The viability plots show that the death rate of the washed early cultures (those who cannot adapt) is higher than the one of the other two for all the replicates of the experiment.

**FIG. S5:**
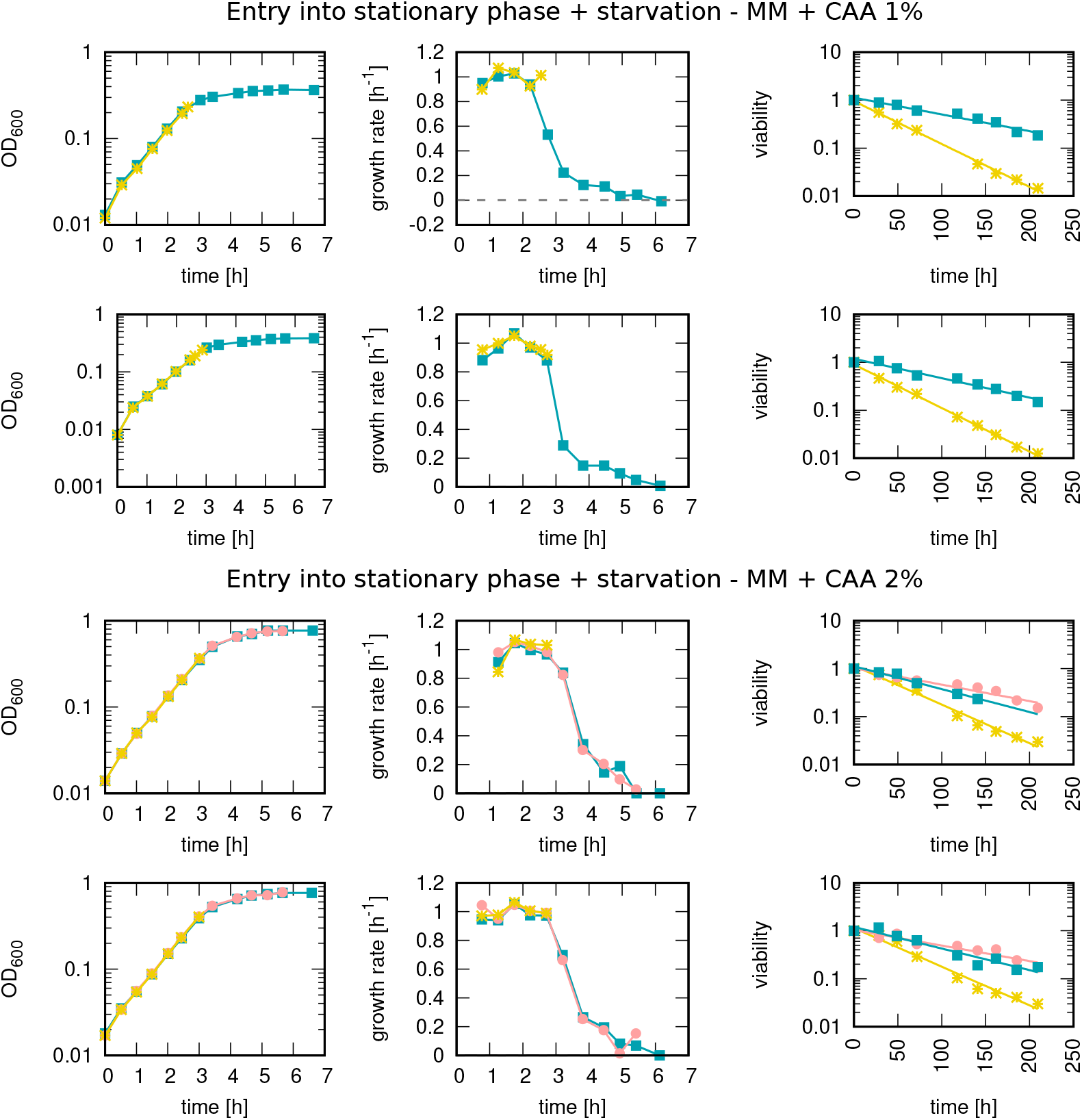
Measurements of OD_600_, growth rate and starvation survival for 2 independent replicates of the experiment with minimal medium plus either 1% or 2% casaminoacids. Yellow asterisks represent data for “washed early” cultures, pink circles for “washed late” cultures, and blue squares for “unwashed” cultures. For each row, the first plot on the left shows the time course of the mass of the colony, measured via the OD_600_, which is shown on the y-axis. The plot in the middle shows the time course of the instantaneous growth rate, with the growth rate on the y-axis and the time on the x-one. The growth rate starts constant during steady-state exponential growth and it drops to zero during the entry into the stationary phase. The growth rate is computed in the same way as in fig S2. The plot on the right shows the survival dynamics of the starved cultures.

**FIG. S6:**
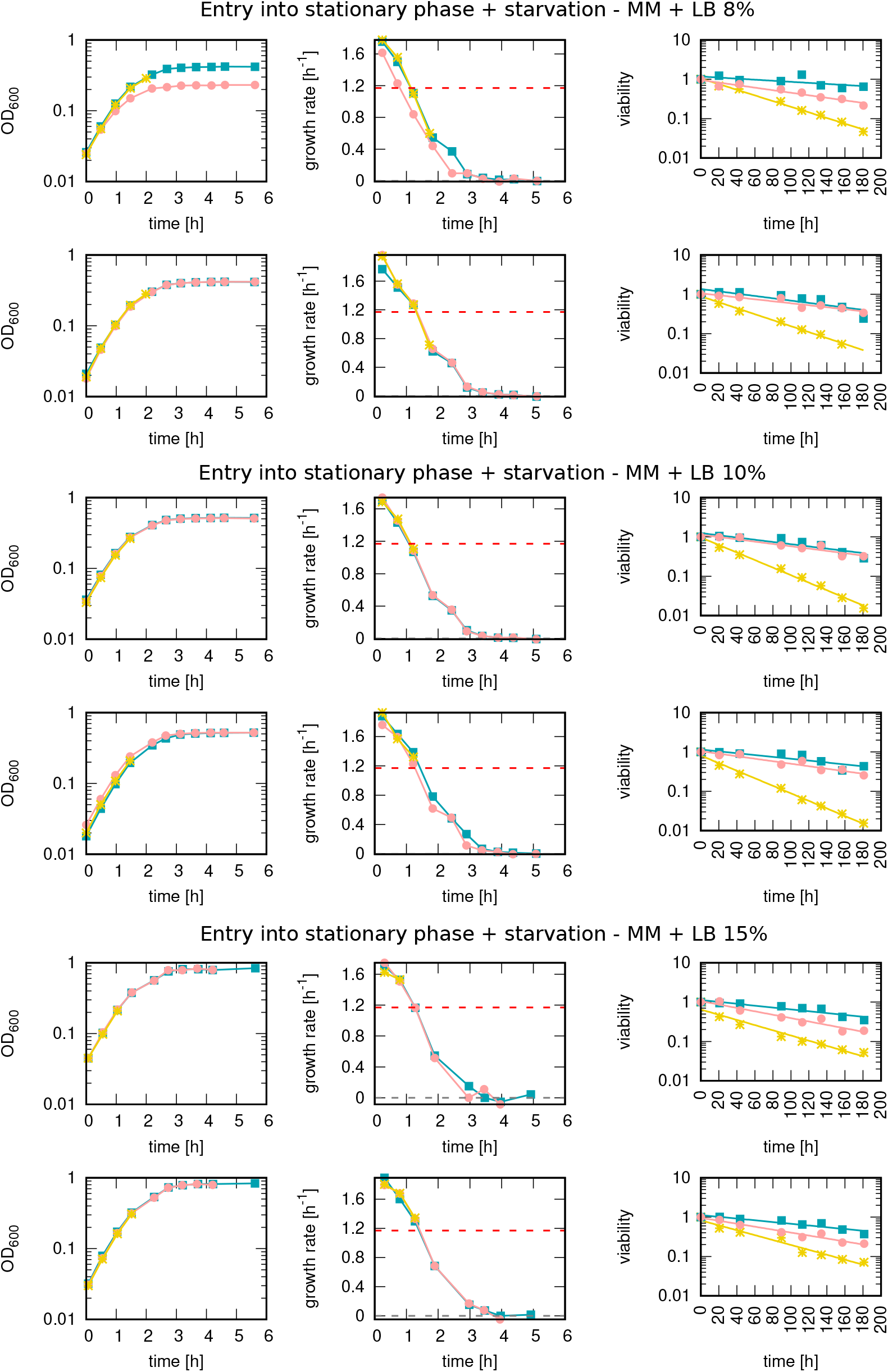
Measurements of OD_600_, growth rate and starvation survival for the 2 independent replicates of the experiment with minimal medium + LB 8%, 10% and 15%. Measurements of OD_600_, growth rate, and starvation survival for the 2 independent replicates of the experiment with minimal medium + LB 8%, 10%, and 15%. Like in fig. S2, each row represents an independent replicate of the experiment with the three cultures: “washed early” (yellow asterisks) “washed late” (pink circles), and “unwashed” (blue squares). For each row, the first plot on the left shows the time course of the mass of the colony, measured via the OD_600_, which is shown on the y-axis. The plot in the middle shows the time course of the instantaneous growth rate, with the growth rate on the y-axis and the time on the x-one. The growth rate is computed in the same way as in fig S2. The growth rate plots are different from the ones in the other experiments, as they show that in this case, the growth rate does not display a phase where is constant, instead it starts from a very high value and starts decreasing right from the start. Therefore, our procedure fails to give a phase of steady-state exponential growth using this medium. The plot on the right shows the survival dynamics of the starved cultures.

**FIG. S7:**
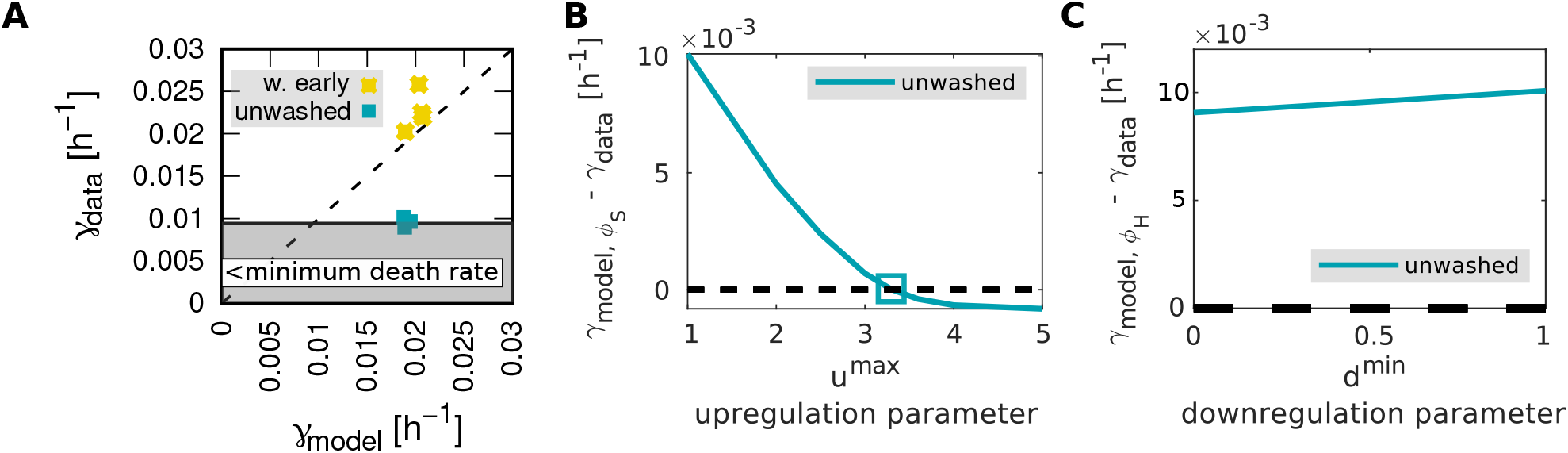
Comparison between the experimental data with the global and targeted regulation models, for the experiment performed with MM + glucose 5%. **(A)** Comparison of the experimental results with the global regulation model predictions. The y coordinate of each point represents the measured death rate, while the x coordinate is the death rate predicted by the global regulation model. The data points are for the washed early (yellow stars) and unwashed cultures (blueish squares). The dashed line is the bisector *γ*_*measured*_ = *γ*_*expected*_, while the shaded area represents the region where the measured death rate is lower than the minimum possible one 0.21day^−1^ [2]. This plot confirms the exponential relation between growth and death rate for the washed culture but also shows that the global regulation hypothesis cannot explain the observed death rate of the two adapted cultures. **(B)** Fit of the value for the up-regulation parameter *u*^max^. The plot shows the results of the targeted regulation model, for different values of the up-regulation parameter. In the simulations, we used the parameters of the unwashed cultures. The y-axis shows the difference between the death rate predicted by the simulation and the measured one, for different values of the x-coordinate *u*^max^. For *u* = 1 the model is identical to the global model. The value for *u* that explains the data on the death rate is given by the intercept between the two solid lines with the dashed line *y* = 0. The plot shows that for both the washed late and unwashed cultures the up-regulation of the good proteome is relatively low, as in the lactose case. **(C)** Fit of the value for the down-regulation parameter *d*^min^. The plot shows the results of the simulation of the targeted regulation model, for different values of the down-regulation parameter. In the simulations, we used the parameters of the unwashed cultures. The y-axis shows the difference between the death rate predicted by the simulation and the measured one, for different values of the x-coordinate *d*. For *d* = 1 the model is identical to the global model. The plot shows that this dynamics is not able to explain the data, as the difference between the measured and the predicted death rate is always greater than zero even in the extreme case of *d* = 0, which corresponds to the pure dilution of the harmful proteome.

**FIG. S8:**
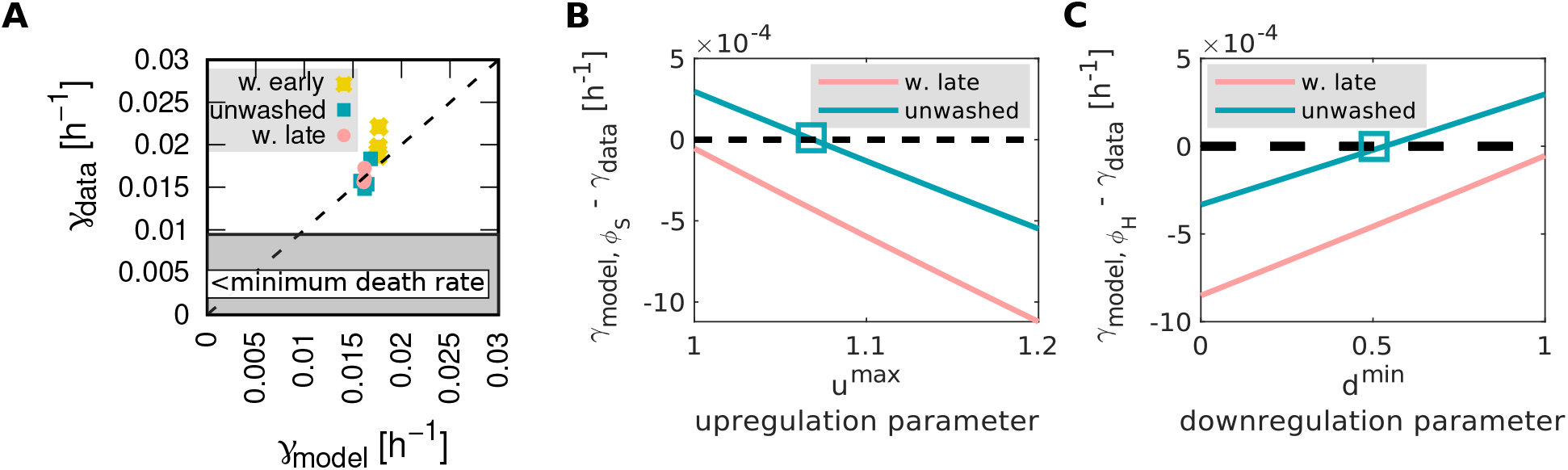
Comparison between the experimental data with the global and targeted regulation models, for the experiment performed with MM + glycerol 5 mM. **(A)** Comparison of the experimental results with the global regulation model predictions. The y coordinate of each point represents the measured death rate, while the x coordinate is the death rate predicted by the global regulation model. The data points are for the washed early (yellow asterisks), washed late (pink rounds), and unwashed cultures (blueish squares). The dashed line is the bisector *γ*_*measured*_ = *γ*_*expected*_, while the shaded area represents the region where the measured death rate is lower than the minimum possible one 0.21day^−1^ [2]. This plot confirms the exponential relation between growth and death rate for the washed culture. Even though there is a difference between the death rates of the adapted and non-adapted cultures, in this case, the adaptation is so low that the global regulation model is able to capture it. This is a consequence of the low growth rate achievable on glycerol. **(B)** Fit of the value for the up-regulation parameter *u*^max^. The plot shows the results of the targeted regulation model, for different values of the up-regulation parameter. In the simulations, we used the parameters of the washed late and unwashed cultures. The y-axis shows the difference between the death rate predicted by the simulation and the measured one, for different values of the x-coordinate *u*^max^. For *u* = 1 the model is identical to the global model. The value for *u* that explains the data on the death rate is given by the intercept between the two solid lines with the dashed line *y* = 0. The plot shows that for both the washed late and unwashed cultures the up-regulation of the good proteome is really close to 1, pointing out again that the global model is sufficient to explain this result. **(C)** Fit of the value for the down-regulation parameter *d*^min^. The plot shows the results of the simulation of the targeted regulation model, for different values of the down-regulation parameter. In the simulations, we used the parameters of the unwashed cultures. The y-axis shows the difference between the death rate predicted by the simulation and the measured one, for different values of the x-coordinate *d*. For *d* = 1 the model is identical to the global model. In this there exists a value of *d*^min^ that can explain the data, *d*^min^ ∼ 0.5, however, this is due to the fact that the global model is sufficient to explain this result, and adding a new parameter just improves the quality of the fit.

**FIG. S9:**
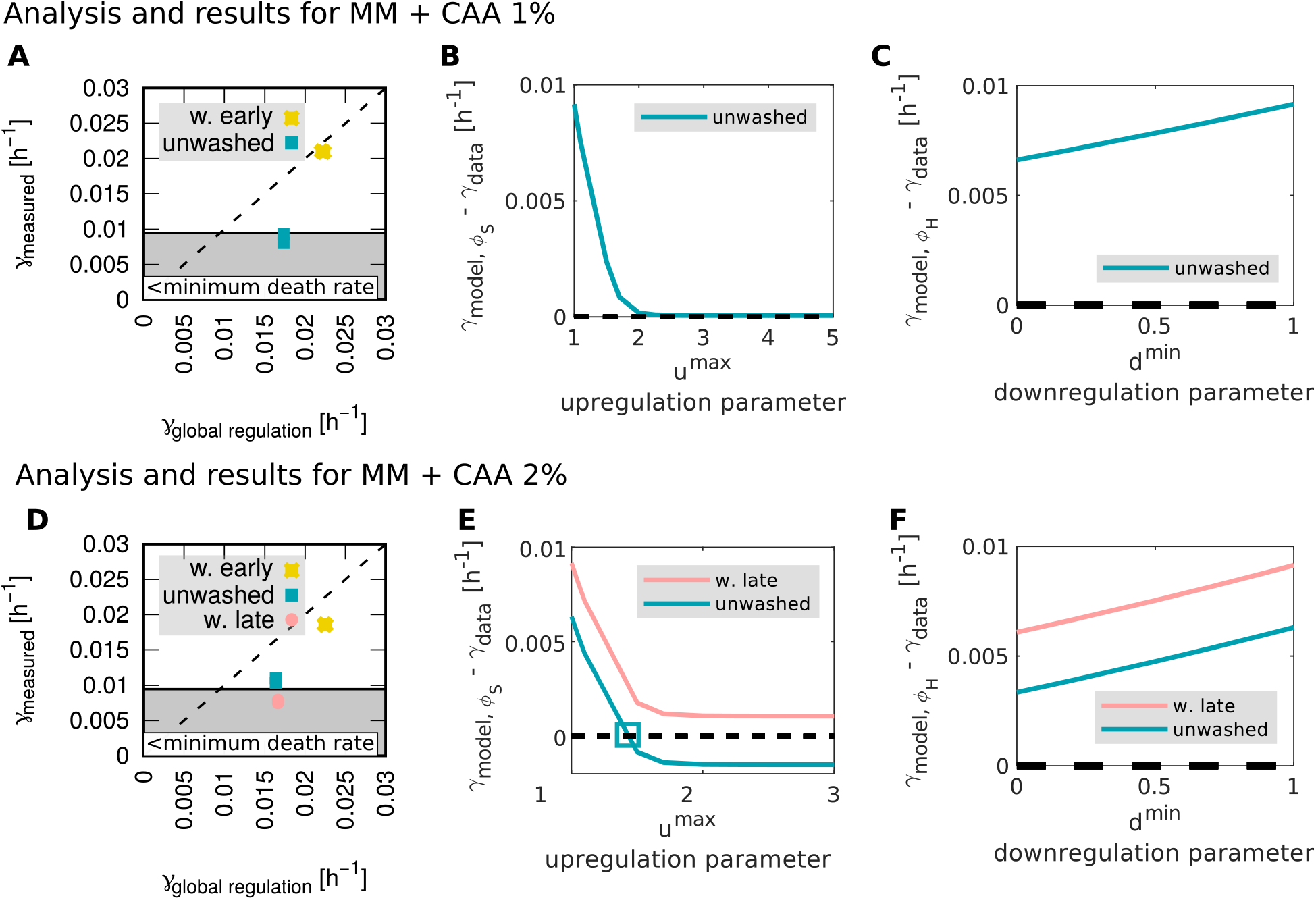
Comparison between the experimental data with the global and targeted regulation models, for the experiment performed with MM + CAA 1%, and 2%. **(A-D)** Comparison of the experimental results with the global regulation model predictions. The y coordinate of each point represents the measured death rate, while the x coordinate is the death rate predicted by the global regulation model. The data points are for the washed early (yellow asterisks) washed late (pink rounds) and unwashed cultures (blueish squares). The dashed line is the bisector *γ*_*measured*_ = *γ*_*expected*_, while the shaded area represents the region where the measured death rate is lower than the minimum possible one 0.21day^−1^ [2]. This plot confirms the exponential relation between growth and death rate for the washed culture, but also shows that the global regulation hypothesis cannot explain the observed death rate of the two adapted cultures. **(B-E)** Fit of the value for the up-regulation parameter *u*^max^. The plot shows the results of the targeted regulation model, for different values of the up-regulation parameter. In the simulations, we used the parameters of the unwashed cultures. The y-axis shows the difference between the death rate predicted by the simulation and the measured one, for different values of the x-coordinate *u*^max^. For *u* = 1 the model is identical to the global model. The value for *u* that explains the data on the death rate is given by the intercept between the two solid lines with the dashed line *y* = 0. The plot shows that the up-regulation of the good proteome is relatively low, as in the lactose case. The lines without an intercept correspond to cultures whose death rate is lower than *γ*_0_, therefore they cannot be explained by either of our models. **(C-F)** Fit of the value for the down-regulation parameter *d*^min^. The plot shows the results of the simulation of the targeted regulation model, for different values of the down-regulation parameter. In the simulations, we used the parameters of the unwashed cultures. The y-axis shows the difference between the death rate predicted by the simulation and the measured one, for different values of the x-coordinate *d*. For *d* = 1 the model is identical to the global model. The plot shows that this dynamics is not able to explain the data, as the difference between the measured and the predicted death rate is always greater than zero even in the extreme case of *d* = 0, which corresponds to the pure dilution of the harmful proteome.

**FIG. S10:**
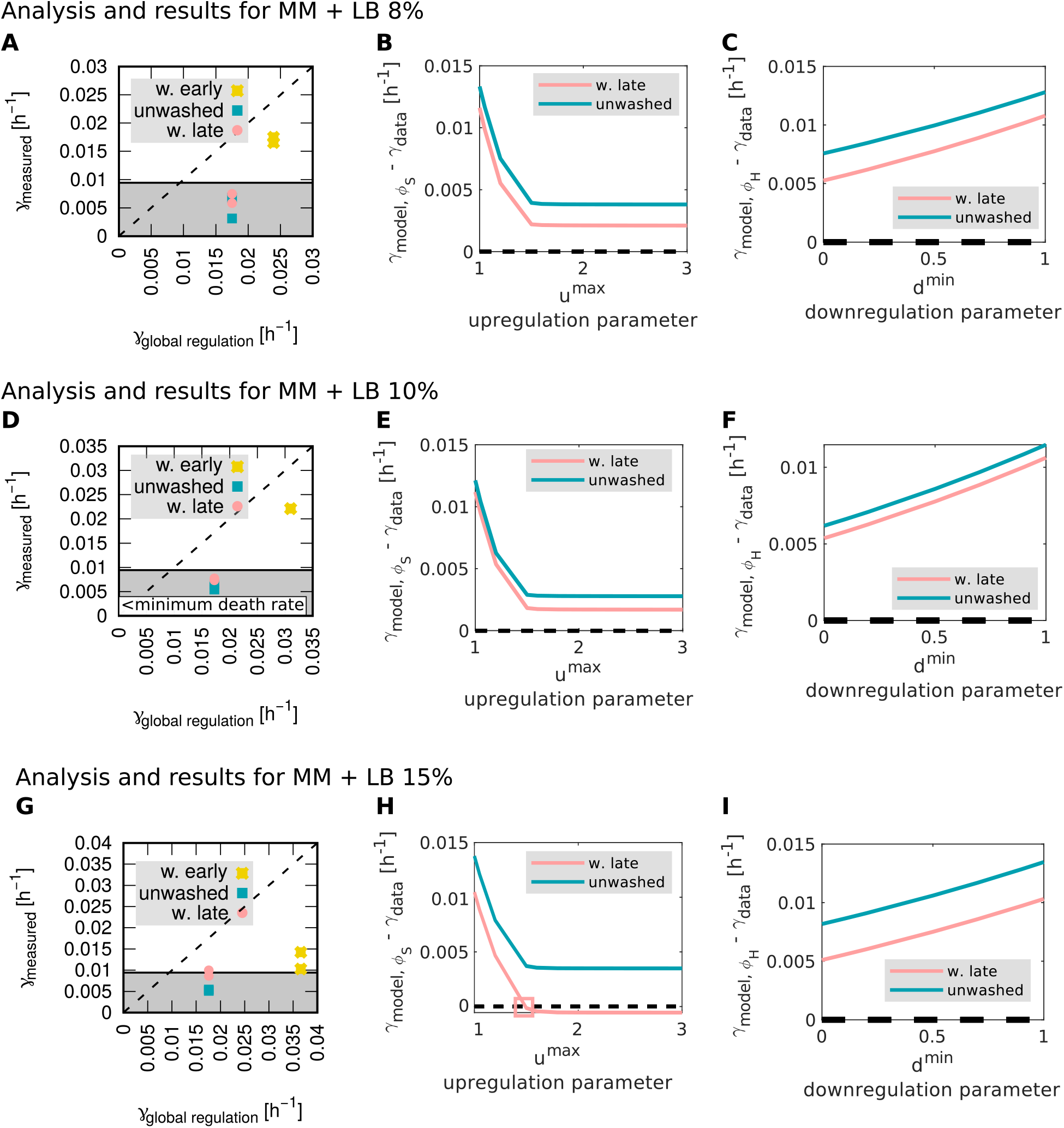
Comparison between the experimental data with the global and targeted regulation models, for the experiment performed with MM + LB 8%, 10%, and 15%. Comparison between the experimental data with the global and targeted regulation models, for the experiment performed with MM + LB 8%, 10%, and 15%. **(A-D-G)** Comparison of the experimental results with the global regulation model predictions. The y coordinate of each point represents the measured death rate, while the x coordinate is the death rate predicted by the global regulation model. The data points are for the washed early (yellow asterisks) washed late (pink rounds) and unwashed cultures (blueish squares). The dashed line is the bisector *γ*_*measured*_ = *γ*_*expected*_, while the shaded area represents the region where the measured death rate is lower than the minimum possible one 0.21day^−1^ [2]. The plots confirm that the adapted cultures die slower, but also show that in this case, we are not able to reproduce the steady-state relation between growth and death rate reported by ref. [2]. This is a consequence of the absence of steady-state growth on this substrate, as shown in fig. S6. **(B-E-H)** Fit of the value for the up-regulation parameter *u*^max^. The plot shows the results of the targeted regulation model, for different values of the up-regulation parameter. In the simulations, we used the parameters of the unwashed cultures. The y-axis shows the difference between the death rate predicted by the simulation and the measured one, for different values of the x-coordinate *u*^max^. For *u* = 1 the model is identical to the global model. The value for *u* that explains the data on the death rate is given by the intercept between the two solid lines with the dashed line *y* = 0. The plot shows that the up-regulation of the good proteome is relatively low, as in the lactose case. The lines without an intercept correspond to cultures whose death rate is lower than *γ*_0_, therefore they cannot be explained by either of our models. **(C-F-I)** Fit of the value for the down-regulation parameter *d*^min^. The plot shows the results of the simulation of the targeted regulation model, for different values of the down-regulation parameter. In the simulations, we used the parameters of the unwashed cultures. The y-axis shows the difference between the death rate predicted by the simulation and the measured one, for different values of the x-coordinate *d*. For *d* = 1 the model is identical to the global model. The plot shows that this dynamics is not able to explain the data, as the difference between the measured and the predicted death rate is always greater than zero even in the extreme case of *d* = 0, which corresponds to the pure dilution of the harmful proteome.

**FIG. S11:**
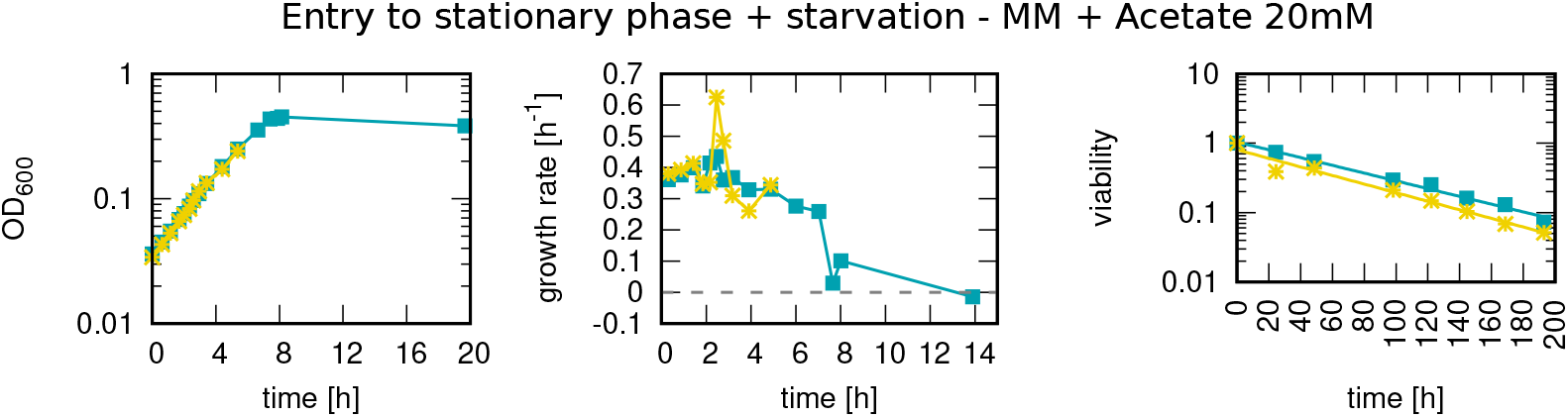
Measurements of OD_600_, growth rate and starvation survival for one replicate of the experiment with minimal medium + acetate 20 mM. Like in fig. S2, each row represents an independent replicate of the experiment with the three cultures: “washed early” (yellow asterisks) and “unwashed” (blueish squares). For this substrate, we do not have the “washed late” culture because at that stage we did not include the second washing step in the procedure yet. For each row, the first plot on the left shows the time course of the mass of the colony, measured via the OD_600_, which is shown on the y-axis. The plot in the middle shows the time course of the instantaneous growth rate, with the growth rate on the y-axis and the time on the x-one. The growth rate starts constant during steady-state exponential growth and it drops to zero during the entry into the stationary phase. The growth rate is computed in the same way as in fig S2. The plot on the right shows the survival dynamics of the starved cultures. The viability plots show that the death rate of the washed early cultures (those who cannot adapt) is almost the same as the one of the other two for all the replicates of the experiment. We think that, as in the case of glycerol, this is due to the very low growth rate achievable on acetate, which caps the adaptation extent and produces a death rate that is not distinguishable.

**FIG. S12:**
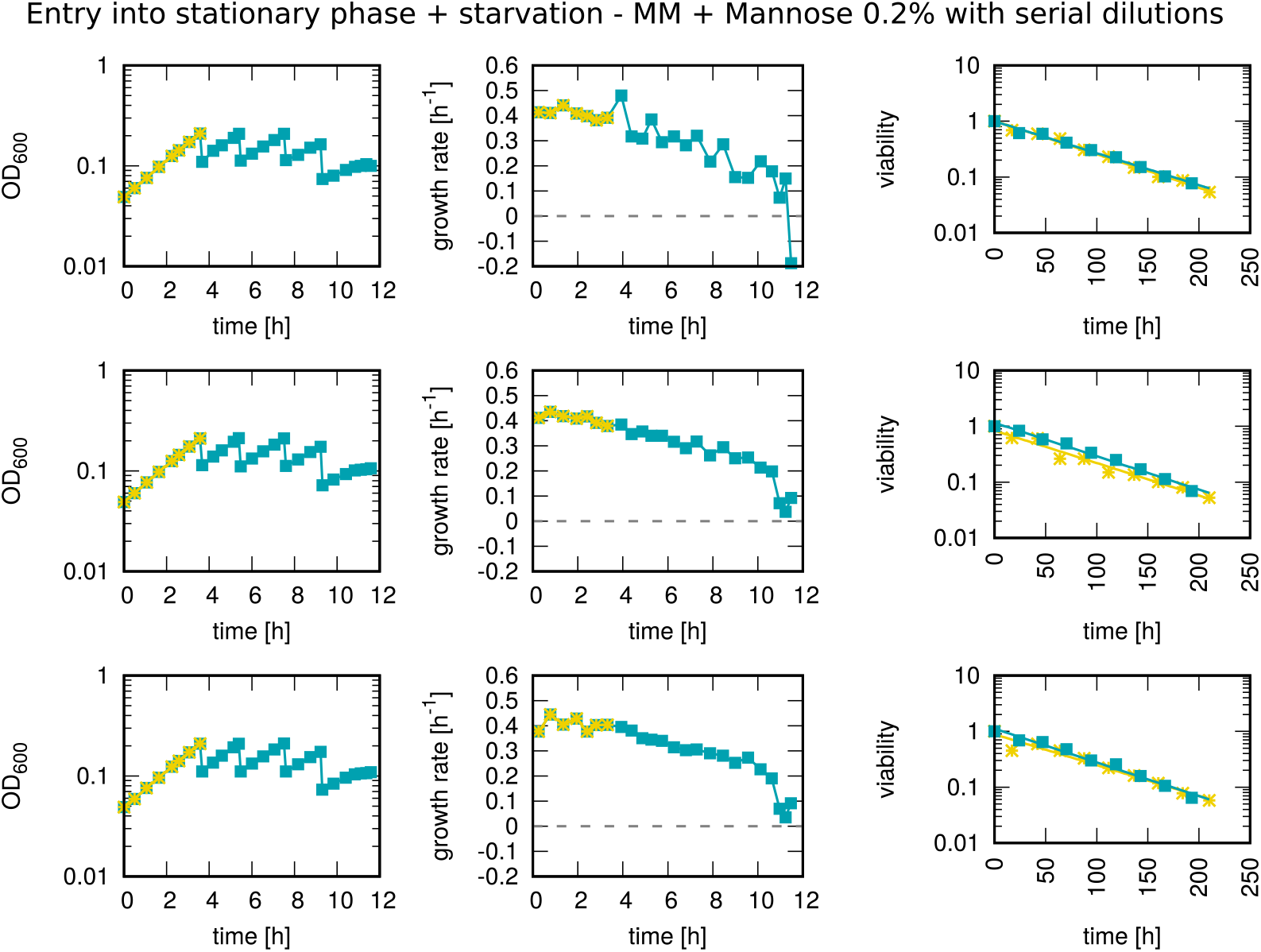
Measurements of OD_600_, growth rate and starvation survival for 3 independent replicates of the serial dilutions experiment with minimal medium + mannose 0.2%. Like in fig. S2, each row represents an independent replicate of the experiment with the three cultures: “washed early” (yellow asterisks) and “unwashed” (blueish squares). For this substrate, we do not have the “washed late” culture because at that stage we did not include the second washing step in the procedure yet. For each row, the first plot on the left shows the time course of the mass of the colony, measured via the OD_600_, which is shown on the y-axis. In this experiment, we performed a dilution every time the OD_600_ reaches a value of 0.2, every time halving the concentration of the nutrient and of the culture (0.2% is the starting concentration). The plot in the middle shows the time course of the instantaneous growth rate, with the growth rate on the y-axis and the time on the x-one. The growth rate starts constant during steady-state exponential growth and it drops to zero during the entry into the stationary phase. The growth rate is computed in the same way as in fig S2. The plot on the right shows the survival dynamics of the starved cultures. The viability plots show that the death rate of the washed early cultures (those who cannot adapt) is almost the same as the one of the other two for all the replicates of the experiment. We think that, as in the case of glycerol, this is due to the very low growth rate achievable on mannose, which caps the adaptation extent and produces a death rate that is not distinguishable.

**FIG. S13:**
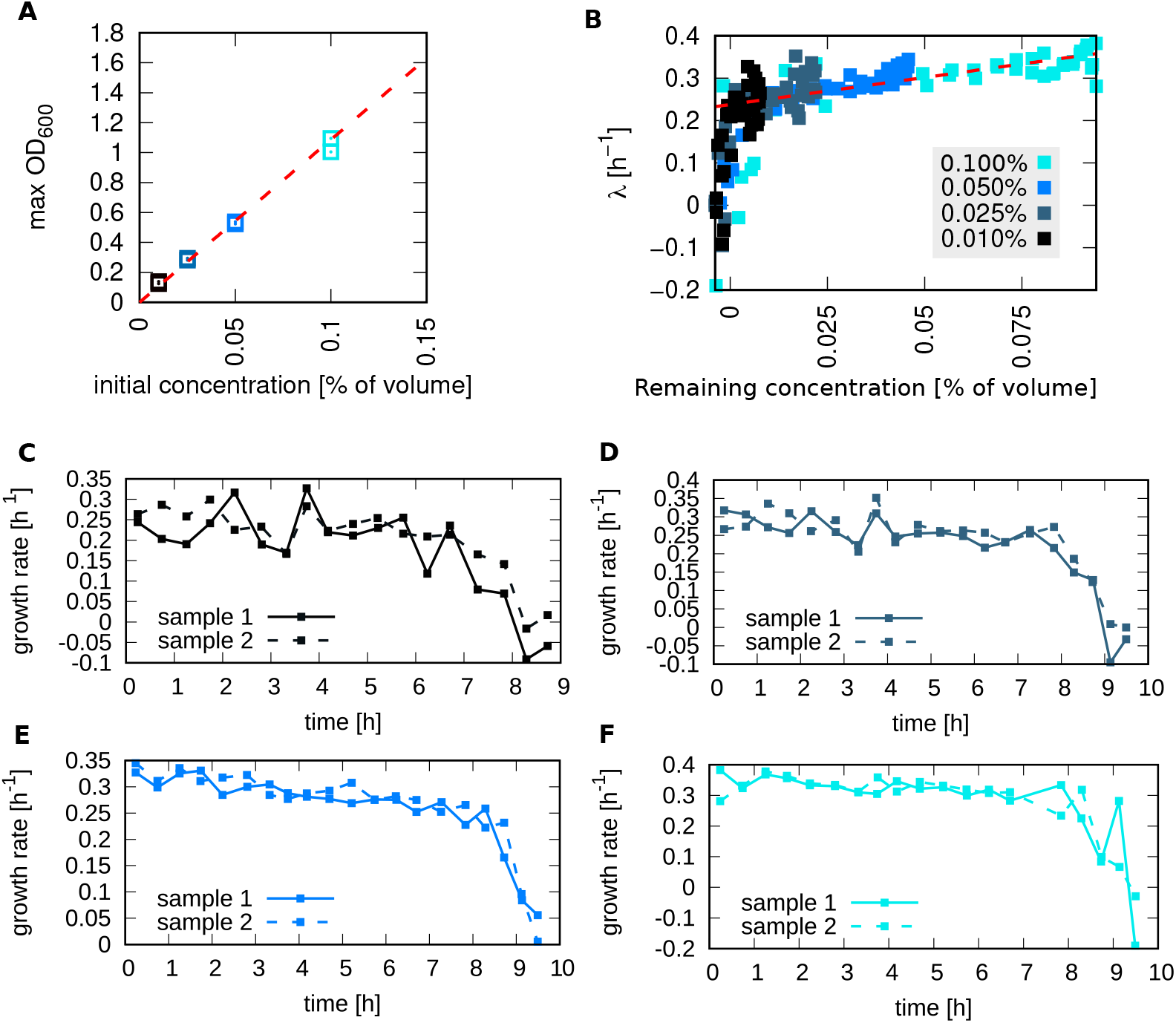
The reasoning behind the design of the dilution experiment. **(A)** The maximum values for the dry mass density linearly depend on the mannose concentration. The horizontal axis shows the initial concentration of mannose, i.e. the total amount of substrate given to the cells, as a percentage of the total volume. The vertical axis shows the maximum reached dry mass density, measured with the optical density OD_600_, for a culture growing on a certain concentration of mannose. The dashed line is a linear fit. The slope of the dashed line is the yield on mannose, equal to 10.8667 OD_600_ *· c*^−1^_Mann._. This quantity allows us to make a correspondence between the dry mass density of a culture and the amount of substrate (per unit volume) that has been consumed by the culture in order to reach that value of OD_600_. **(B)** The instantaneous growth rate of a culture growing on mannose depends on the remaining concentration of the substrate. Each point corresponds to a growth rate measurement. For each point, the y-coordinate is the growth rate measured at a certain time t during the growth experiment. The x-coordinate is the remaining concentration of substrate, computed using the optical density OD_600_ of the culture at time t and the value for the yield. The figure present eight data series, two for each of the tested concentrations listed in the legend. The plot shows that for medium to high substrate concentration, the growth rate depends linearly on it, and the red curve is a fit of this behavior. Instead, when the concentration is approaching zero the growth rate shows a sharp jump. **(C-F)** Time course of the growth rate for growth on different mannose concentrations. For each tested concentration we performed two replicates of the experiment. The horizontal axis shows the time measured in hours, while the vertical one shows the growth rate of the two cultures. These panels show that even if the initial concentration of the nutrient changes the cells need roughly the same amount of time to deplete it, around 9h.

**FIG. S14:**
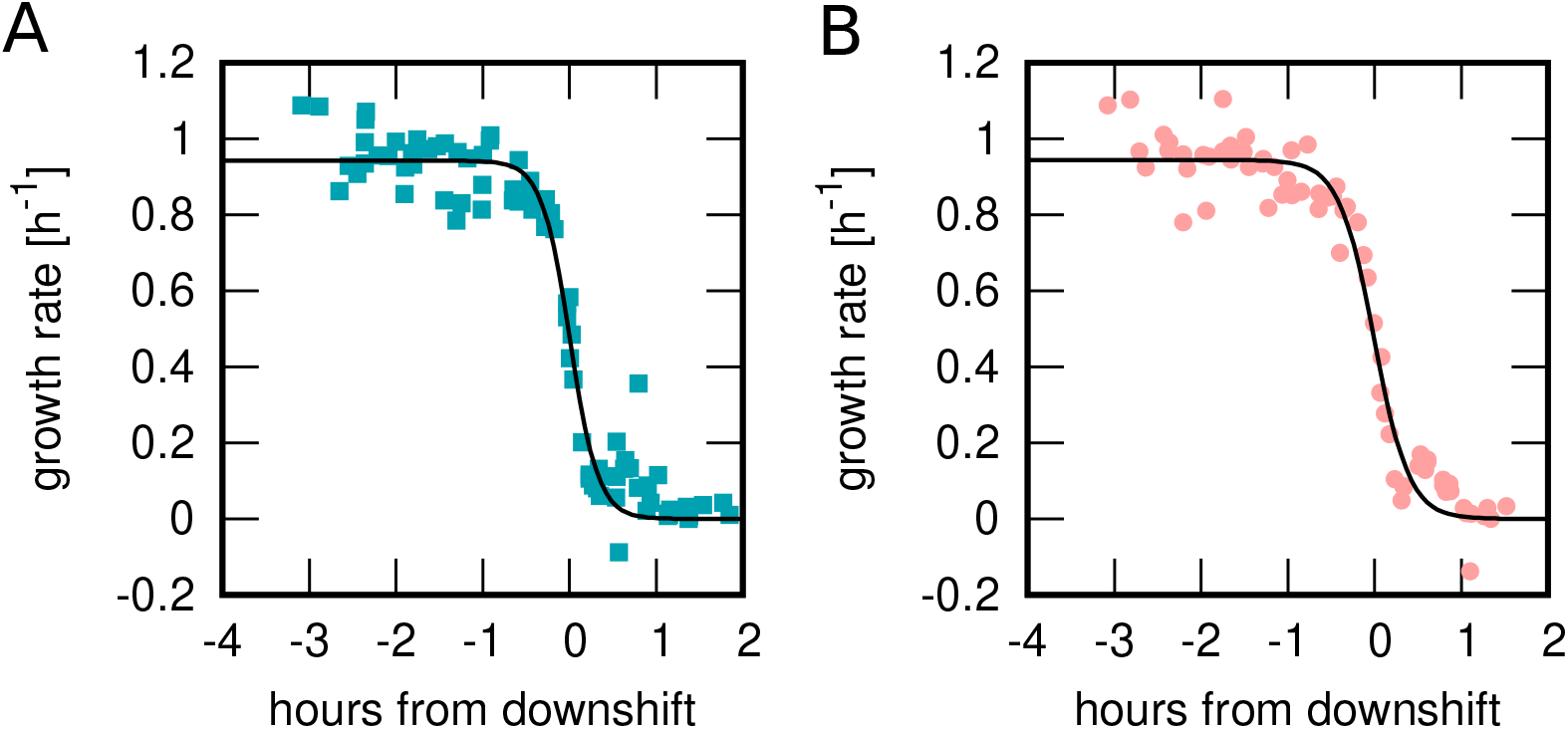
Sigmoid functions can capture the dynamics of the growth rate during the entry into the stationary phase. Example of the sigmoidal fit of the growth rate, for the 7 replicates of the experiment with minimal medium + lactose 0.05%. Data and fit are for the unwashed (panel A) and washed late cultures (panel B). The panels show that the sigmoidal fit can capture both the first and second plateau, corresponding to *λ*^*^ and *λ* = 0, and the entry into starvation phase in between.

**FIG. S15:**
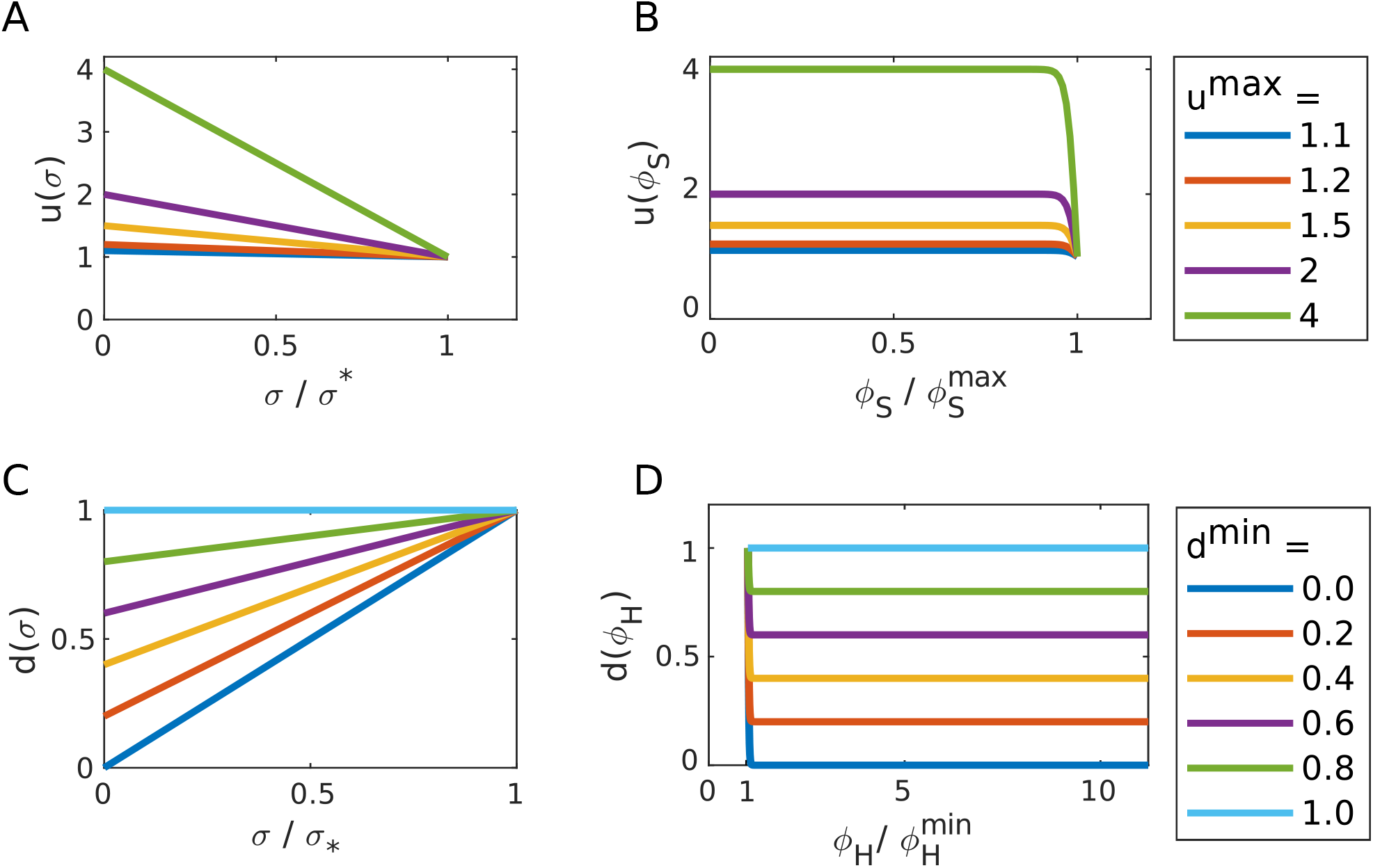
Dependence on translational activity *σ* and sector size *ϕ*_H_ and *ϕ*_S_ of the up- and down-regulation parameters *u*(*t*) and *d*(*t*). Dependence on translational activity *σ* and sector size *ϕ*_H_ and *ϕ*_S_ of the up- and down-regulation parameters *u*(*t*) and *d*(*t*). As eq.s 21 and 22 of the main text prescribe, these two parameters activate (i.e. they change from their steady-state value = 1) during the entry into the stationary phase, and they remain active until *ϕ*_*S*_ and *ϕ*_*H*_ have reached their extreme values. We modeled these two behaviors by defining *u*(*t*) and *d*(*t*) as

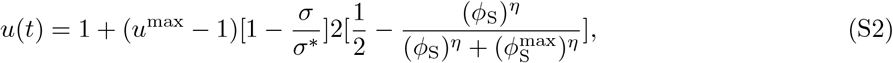

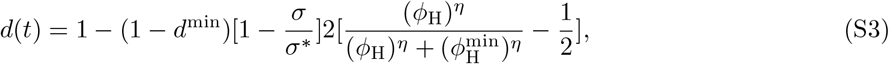

equations 32 and 33 of the main text. Panels A-C show the dependence of the parameters on the translational activity *σ*, they show that *u*(*t*) and *d*(*t*) increase/ decrease linearly towards *u*^max^ and *d*^min^ as *σ* start to decrease respect to the steady state value *σ*^*^. Panels B-D show the dependence on the harmful and survival sector size. *u*(*t*) and *d*(*t*) stay constant while 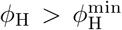 and 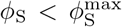, and they deactivate (i.e. they return to 1) otherwise.

